# Integrative modelling reveals the structure of the human Mic60-Mic19 subcomplex and its role as a diffusion barrier in mitochondria

**DOI:** 10.64898/2026.01.30.702776

**Authors:** Evangelia Nathanail, Edoardo Rolando, Max Ruwolt, Iryna Zaporozhets, Fan Liu, Cecilia Clementi, Oliver Daumke

## Abstract

Mitochondrial crista junctions (CJs) operate as regulated gateways into the cristae microenvironment, whose protein, metabolite, and ion compositions are finely tuned for mitochondrial function. The Mic60-Mic19 complex of the mitochondrial contact site and cristae organizing system (MICOS) complex was suggested to span across CJs and act as a diffusion barrier, but little is known of how its dynamic architecture facilitates this task. To address this open question, we determined the crystal structure of an amino-terminal dimeric helical bundle of human Mic60. These and previous structural and biochemical data were harnessed in molecular dynamic (MD) simulations to develop a dynamic model of the human tetrameric Mic60-Mic19 subcomplex in the CJ environment, to validate its architecture using *in organello* cross-linking data and to computationally characterize its function as a diffusion barrier. Our integrative structural biology approach enables the functional investigation of flexible, multidomain protein complexes which escape conventional structural biology methods.

## Introduction

Mitochondria are highly dynamic organelles that serve as central hubs for energy production, amino acid and fatty acid metabolism, and the regulation of apoptosis in eukaryotic cells (Kondadi & Reichert, 2024; Suomalainen & Nunnari, 2024; Tábara et al., 2025). The morphology and plasticity of their network vary significantly across different organisms, tissues, and even within individual cells, reflecting the intricate relationship between their diverse cellular roles and their structure (Daumke & van der Laan, 2025; Quintana-Cabrera & Scorrano, 2023; Ryu et al., 2024). Mitochondrial networks are characterized by their double membrane architecture (Sjostrand, 1953), comprised of a semi-permeable outer mitochondrial membrane (OMM) and an impermeable inner mitochondrial membrane (IMM). The IMM features a vastly extended surface area due to cristal invaginations that extend from the intermembrane space (IMS) into the matrix.

Cristae function as specialized microenvironments within the IMS and the IMM, where the oxidative phosphorylation machinery is located (Davies et al., 2011; Wurm & Jakobs, 2006). These regions are characterized by distinct protein and lipid compositions that coordinate the intricate cristal membrane architecture and optimize mitochondrial adenosine triphosphate (ATP) production (Ikon & Ryan, 2017; Mühleip et al., 2023; Venkatraman et al., 2023; Zheng et al., 2024). ATP synthase dimer ribbons stabilize negative curvature at the cristal rim (Strauss et al., 2008), while Optic Atrophy 1 (OPA1) forms ring-like assemblies essential for maintaining the morphology of the cristal lumen (reviewed in Daumke & van der Laan, 2025; Faelber et al., 2019; Nyenhuis et al., 2023; von der Malsburg et al., 2023). A critical architectural component of cristae for establishing a distinct mitochondrial microenvironment is the crista junction (CJ), which acts as a diffusion barrier between cristae and the IMS, regulating the passage of proteins, metabolites, and ions (Frey & Mannella, 2000; Rampelt et al., 2017).

The mitochondrial contact site and cristae organizing system (MICOS) complex localizes to CJs and plays a crucial role in their formation and stability (Harner et al., 2011; Hoppins et al., 2011; von der Malsburg et al., 2011). MICOS is a conserved multi-subunit protein complex of approximately 700 kDa in size and can be biochemically separated into the Mic10 subcomplex (containing Mic10, Mic26 and Mic27) and the Mic60 subcomplex (containing Mic60, Mic19 and its paralog Mic25 in vertebrates). The Mic12 subunit serves as a bridging component between the subcomplexes in yeast (Anand et al., 2016; Guarani et al., 2015; John et al., 2005; Mun et al., 2010), although this was recently disputed for its putative mammalian orthologue Mic13 (Naha et al., 2024). The Mic10 subcomplex is predominantly membrane-embedded and is thought to form a curved scaffold essential for stabilizing CJs (Barbot et al., 2015; Bohnert et al., 2015; Rampelt et al., 2022; Stephan et al., 2024). In contrast, the Mic60 subcomplex is largely exposed to the IMS and anchored to the IMM by a single N-terminal transmembrane helix (TM) of Mic60 (van der Laan et al., 2016). Deletion of either Mic10 or Mic60 results in a near complete loss of CJs, with cristae forming lamellar sheet-like structures that are disconnected from the IMS (Harner et al., 2011; Hoppins et al., 2011; von der Malsburg et al., 2011). In addition to shaping the IMM, the Mic60 subcomplex interacts with proteins of the OMM, such as the sorting and assembly machinery (SAM), bridging the intermembrane space and contributing to mitochondrial protein import and trafficking (Korner et al., 2012; Tang et al., 2020; Utsumi et al., 2018; Xie et al., 2007; Zerbes et al., 2012).

Though MICOS is essential for regulating mitochondrial form and function, the molecular mechanism by which it stabilizes CJs remains poorly understood. First structural and biochemical insights on the Mic60 subcomplex were derived from thermophilic yeast orthologues. They revealed that Mic60 exists in a dimeric, autoinhibited state (Hessenberger et al., 2017), while Mic19 binding induces Mic60 tetramerization via a conserved interface in the coiled-coil (CC) domain (Bock-Bierbaum et al., 2022). The C-terminal mitofilin domain forms a domain-swapped dimer featuring a bent peripheral membrane binding site, which may bind to the highly curved CJ membrane. The Mic60-Mic19 subcomplex was suggested to span across the CJ and act as a mechanical strut preserving CJ diameter (Bock-Bierbaum et al., 2022).

While the current Mic60-Mic19 model was derived from fungal MICOS components, comparable data on the Mic60 subcomplex of higher eukaryotes have remained scarce. However, such data are crucial to understand the function of the Mic60-Mic19 complex in human mitochondria, particularly since both CJ morphology and Mic60 size vary between animals and fungi (Collins et al., 2002; Zick et al., 2009). The lack of structural data for the human complex may be attributed to Mic60’s high structural flexibility and the intricate oligomerization network of the Mic60-Mic19 complex, which hinders structural characterization by conventional structural biology techniques. Recent efforts sought to investigate the structure of the subcomplex by AlphaFold predictions (Jumper et al., 2021) and to further validate these models through a subset of cross-links from human mitochondria (Bartolec et al., 2023) or to characterize them in large-scale molecular dynamics simulations of mitochondrial cristae (Brown et al., 2025). However, AlphaFold predictions of homo-oligomeric coiled-coils, such as the core of Mic60, are often problematic (Madaj et al., 2025). In addition, these models did not consider the hetero-octameric stoichiometry of the Mic60-Mic19 complex. Accordingly, the structural predictions remained incomplete and the role of the Mic60-Mic19 subcomplex in the control of CJ architecture and function was not addressed in these studies.

Here, we present an integrative structure biology approach to model the human Mic60-Mic19 complex in its assembled form in a CJ-like environment, combining a novel experimental dimeric crystal structure of human Mic60 with *in vitro*, *in silico*, and *in organello* data. Experimentally solved and biochemically validated structures of nearly all folded regions of Mic60 across multiple organisms provide the foundation for this model, which was further refined through molecular dynamics simulations and validated by cross-linking mass spectrometry of intact mitochondria. The high dynamics of the Mic60-Mic19 complex facilitates its function as a diffusion barrier, as analysed by coarse-grained molecular dynamics simulations. Our approach highlights the pivotal and multifaceted role of the Mic60-Mic19 subcomplex in maintaining CJ architecture and function and serves as an exemplary pipeline for studying dynamic multi-domain protein assemblies which have so far escaped structural characterization due to limitations of conventional structural biology methods.

## Results

### Human Mic60 features an N-terminal dimeric helical bundle

While previous structural studies of MICOS were based on fungal Mic60 and Mic19 orthologues (Bock-Bierbaum et al., 2022), structural data on the human MICOS complex have remained limited. Animal Mic60 harbours a predicted helical bundle (HB) situated between the intrinsically disordered region (IDR) and the CC domain, which is not present in the fungal Mic60 counterparts (Figure 1A, Supplementary Figure 1). An evolutionary analysis based on the framework presented by Huynen et al. (2016) indicates that the Mic60 HB has emerged during animal evolution as the organisms acquired more complex multicellularity (Supplementary Figure 1A).

**Figure 1:**
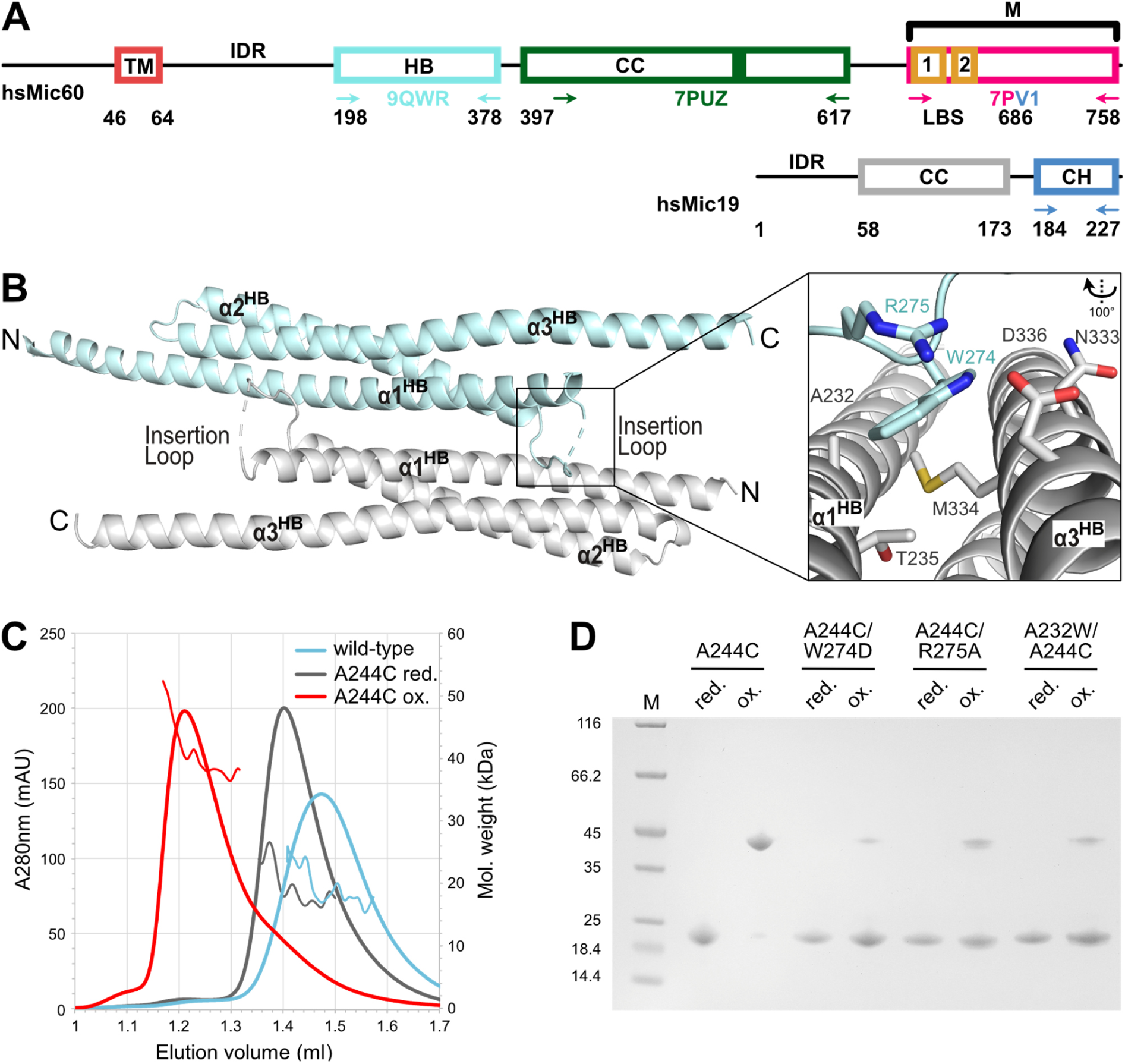
Animal Mic60 orthologues possess an N-terminal helical bundle with a conserved fold. **A** Domain architecture of hsMic60 and hsMic19, with residue numbers annotated below the domains. Solved orthologue crystal structures are indicated by arrows and PDB ID (helical bundle (HB) in cyan (PDB ID: 9QWR), coiled-coil (CC) in dark green (PDB ID: 7PUZ), mitofilin domain in pink and CHCH domain in blue (PDB ID: 7PV1)). **B** Cartoon representation of the hsMic60 N-terminal helical bundle (Mic60 HB). N- and C-termini of each monomer are labelled. Close-up view of the dimerization interface via an antiparallel loop insertion of residue W274, coordinated by R275, into an interface composed of helices α1^HB^ and α3^HB^ of the opposite moiety. **C** Size-exclusion chromatography and Right-Angle Light Scattering (SEC-RALS) profile of wild-type Mic60 HB (cyan) and the A244C variant in a reduced (grey) and oxidized (red) state captures transient dimer formation. **D** Non-reducing SDS PAGE of Mic60 HB cysteine mutants before and after CuSO_4_ oxidation. For quantification, see Supplementary Figure 3C.

We recombinantly expressed the HB domain of human Mic60 (homo sapiens (hs) Mic60, residues 198-378, hereafter referred to as Mic60 HB, Figure 1A) and purified it to homogeneity (Supplementary Figure 2). Crystals of this construct diffracted to 2.78 Å resolution, and the structure was solved by molecular replacement using an AlphaFold prediction of the HB domain as a template (PDB ID: 9QWR, see Supplementary Table 1 for data collection and refinement statistics). The structure of the HB domain comprises an elongated three-helix bundle of helices α1^HB^-α3^HB^ (Figure 1B). It features a hydrophobic, conserved core, pointing to stable domain architecture (Supplementary Figure 1B).

Interestingly, the two HB domain monomers in the asymmetric unit self-associated into a two-fold symmetric antiparallel dimer (Figure 1B). Dimerization is mediated primarily by helix α1^HB^, with the interface spanning an area of 1060 Å^2^, stabilized by a symmetric loop insertion (Supplementary Figure 3A). Residues W274 and R275 of the inserting loop embed into a hydrophobic groove composed of helices α1^HB^ and α3^HB^ of the opposite Mic60 HB monomer. Notably, W274 and R275 are surface-exposed and highly conserved in vertebrate Mic60 (Supplementary Figure 1B, C), pointing to their functional importance.

We characterized the assembly status of the Mic60 HB by size-exclusion chromatography coupled to right angle laser scattering (SEC-RALS), in which the protein eluted as a monomer (Figure 1C). To identify a possible transient dimer, we engineered a structure-based disulfide bond by introducing the A244C mutation in the centre of the hydrophobic dimerization interface (Supplementary Figure 3A). Indeed, Mic60 HB A244C predominantly formed dimers under oxidative conditions (Figure 1C). When the disulfide-stabilizing variant was combined with mutations either within the loop (W274D, R275A) or in the hydrophobic groove of the opposite monomer (A232W), the combinatorial mutants showed markedly reduced dimer formation. These data are in agreement with a model indicating that W274 and R275 mediate transient dimerization of Mic60 HB (Figure 1D, Supplementary Figure 3B, C). Accordingly, the HB domain may act as a novel oligomerization domain in animal Mic60.

Despite low sequence similarity, a FoldSeek (van Kempen et al., 2024) homology search revealed that the Mic60 HB possessed highest structural similarity to the IMS protein Smac/DIABLO, which is a known apoptosis regulator (Chai et al., 2000) (Supplementary Figure 3D). Like Mic60 HB, Smac/DIABLO also forms a dimer; however, it employs a distinct dimerization interface and lacks a structural equivalent to the insertion loop observed in Mic60 HB (Supplementary Figure 3E). These observations indicate an unexpected evolutionary relation of the MICOS complex and apoptosis regulators in animalia.

### Assembling a homology-based structural model of the human Mic60-Mic19 complex

Three crystal structures cover the assembly of nearly all folded regions of Mic60: the dimeric Mic60 HB domain presented here, the tetrameric CC domain of *Lachancea thermotolerans* Mic60 (PDB ID: 7PUZ) and a fusion construct containing the dimeric Mic60 mitofilin domain and the Mic19 CHCH domain from *Chaetomium thermophilum* (PDB ID: 7PV1). Building on these structures, we constructed a homology-based model of the human Mic60-Mic19 hetero-octameric complex. As structural prediction algorithms failed to accurately reproduce the known biochemical interactions, we assembled the complex manually. To this end, human homology models of the tetrameric coiled-coil region and the mitofilin domain were generated using SWISS-MODEL (Waterhouse et al., 2018) and combined with AlphaFold predictions of disordered or loop regions to construct a tetrameric model of human Mic60 (Jumper et al., 2021). The CHCH domain of human Mic19 was positioned on the human Mic60 mitofilin region based on the previously published structure of the *Chaetomium thermophilum* complex, while the remaining Mic19 regions were excluded from the model due to ambiguous conformations in AlphaFold predictions. To characterize the assembly of the human Mic60 CC domain, we employed blue native gel polyacrylamide (BN-PAGE) and SEC-RALS analysis of two CC constructs with different C-terminal truncations. Consistent with results previously obtained for *Lachancea thermotolerans* Mic60 CC, the human Mic60 CC domain appeared in dimeric or tetrameric forms, where the assembly was controlled by its adjacent regions (Supplementary Figure 4A, B).

The assembled human Mic60-Mic19 model aligns with prior biochemical and structural observations and centres around a tetrameric CC core (Figure 2A, box 1). Importantly, the tetrameric CC supports the formation of the two C-terminal dimeric mitofilin-CHCH modules, which were described to form a highly curved membrane-binding region (Figure 2A, box 2) (Bock-Bierbaum et al., 2022). Surrounding the core, the Mic60 HB dimers and the highly flexible IDR occupy the remaining space (Figure 2A, box 3; IDR not shown). Short inter-domain linkers result in a mostly planar configuration of the complex, oriented perpendicularly to the CJ neck.

**Figure 2:**
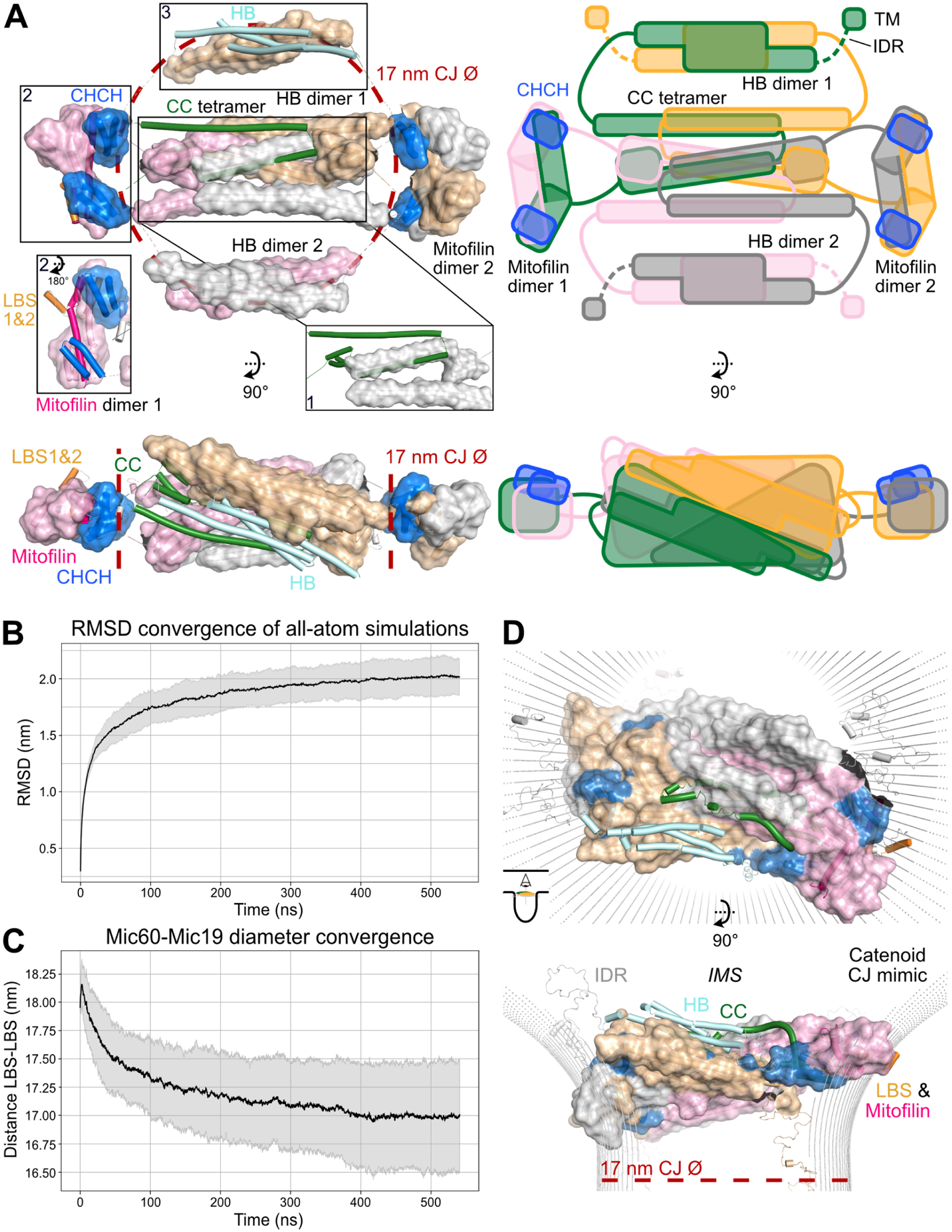
MD simulations of a homology-based structural model of the human Mic60-Mic19 complex on a CJ mimic. **A** Homology-based model of the human Mic60-Mic19 complex based on known crystal structures, displayed as cylinders and surface. Mic60 monomer one is coloured according to annotated domains in Figure 1A, monomers two, three and four in sand, grey and pink, respectively. The IDR is not displayed for illustration purposes. For Mic19, only the CHCH domains could be modelled confidently, shown here in blue. Known oligomerization domains of Mic60 are displayed in boxes 1 (tetrameric CC, here only one dimer displayed for illustration purposes, PDB ID: 7PUZ, monomer 1 in forest green), 2 (mitofilin dimer, PDB ID: 7PV1, monomer 1 in orange and hot pink) and 3 (dimeric Mic60 HB, monomer 1 in cyan). Dashed lines indicate the width of a human CJ (17 nm) and the perpendicular placement of the complex in the CJ. A graphical sketch of the model is presented on the right, with monomer 1 coloured in green and other monomers coloured as in A (sand, grey and pink). **B** Root-Mean-Square deviation (RMSD) average and standard deviation from the 30 all-atom simulations is shown over the time course of the simulation. **C** Diameter convergence of the human Mic60-Mic19 complex during the 30 all-atom simulations, prior to the attachment to the catenoid. **D** Final snapshots of the Cα structure-based MD simulations of the homology-based model represented as cylinders and surface. The base diameter of the catenoid, indicated in a red dashed line, is 17 nm.

Due to missing structural data describing the interactions between the folded domains within the Mic60-Mic19 complex, linkers were initially added in an extended conformation to prevent domain clashes. As a result, the initial homology-based model adopted an overstretched, energetically strained conformation, with a lateral width exceeding the average CJ diameter of 17 nm observed in human mitochondria (Fry et al., 2024; Rabl et al., 2009) (see red dotted line in Figure 2A).

We could not experimentally determine how the HB dimers are positioned relative to the tetrameric CC. An alternative model was therefore generated which was significantly more extended (Supplementary Figure 4C, D). Further *in organello* validation (see below) supported the first, more compact model on which the following computational analyses were focused.

### Molecular dynamics simulations reveal the dynamic conformation of the human Mic60-Mic19 complex

To relax the unfavourable conformation and obtain dynamic information of the Mic60 IDRs in particular, we used the generated homology model as a starting point for molecular dynamics (MD) simulations. Domains were stabilized to reflect their assembly in the crystals, while no constraints were imposed on additional inter-domain contacts or unstructured regions.

Within 500 ns of initial all-atom MD simulations, an equilibration state was reached, as indicated by root-mean-square deviation (RMSD) convergence (Figure 2B, Supplementary Figure 5A). Notably, the simulations led to a compression of the structure along its longitudinal size, ultimately converging to an average width of 17 nm, as measured between the LBS regions (Figure 2C, Supplementary Figure 5B), which is consistent with previously reported diameters of human CJs (Fry et al., 2024; Rabl et al., 2009). Notably, the alternative model retained a higher average equilibrium width (Supplementary Figure 5C, D).

The initial main model was used in parallel to initiate machine-learned coarse-grained (MLCG) simulations in order to explore faster conformational dynamics (Charron et al., 2025).

The MLCG simulations accordingly exhibit faster convergence compared to the all-atom simulations (Supplementary Figure 6A) while retaining a similar average secondary structure content and overall architecture as the all-atom simulations, revealing a consistent fold for the MD model of the Mic60-Mic19 complex (Supplementary Figure 6B).

To characterize the membrane interaction of the Mic60-Mic19 complex, we performed coarse-grained MD simulations of the mitofilin-CHCH dimer using the Martini 3 force field, embedding it in a lipid bilayer with composition and curvature representative of the IMM environment. The mitofilin domain strongly bound to this membrane, with a substantial free energy gain of approximately 100 kJ/mol (Supplementary Figure 7A, B). This supports a model in which the mitofilin domain dimer attaches with high affinity to the IMM.

To integrate the structural model into an environment mimicking the CJ architecture, we attached the relaxed all-atom Mic60-Mic19 models to a catenoid surface with a base diameter of 17 nm, using the first residue following the transmembrane domain (Ile62) and the LBS2 in the mitofilin domain as surface attachment points (Supplementary Figure 7C, D). The membrane interaction strength was modelled to reflect the ∼100 kJ/mol binding free energy observed in the coarse-grained simulations of the mitofilin-CHCH dimer. Although the all-atom model was free to move along the catenoid and, consequently, adapt to its width, it maintained an average width of 17 nm, consistent with the value reached during the prior unconstrained all-atom equilibration (Figure 2C).

Both the all-atom force field and the MLCG model led to an artificial compression of the IDRs (Kasahara et al., 2019) (Supplementary Figure 7C, D). To more accurately capture IDR dynamics, the final frames from each of the 30 all-atom runs were therefore converted into coarse-grained Cα structure-based models (Clementi et al., 2000), leading to a refined final structure (Figure 2D). Overall, the MD simulations revealed a more compact organization of the inter-domain linker regions, with the dimeric Mic60 HB collapsing onto the tetrameric CC on both sides. AlphaFold2 predicted an extended helical connection between the Mic60 HB and the CC (see Materials and Methods), which became sandwiched between these domains during the simulations, forming a more compact core structure that covered almost the entire pore of the CJ. This folded core remained stable throughout the simulation, while in the coarse-grained Cα structure-based simulations, the IDRs were flexible, covering diverse trajectories within the remaining CJ space. In this way, the IDRs formed an extended network above and below the HB-CC core.

To validate our MD simulations and obtain information on the Mic60-Mic19 structure in an organellar context, we utilized a publicly available human mitochondrial cross-linking dataset employing the enrichable and MS-cleavable cross-linker disuccinimidyl bis-sulfoxide (DSBSO) (Zhu et al., 2024). The database contains 236 unique cross-links for hsMic60 distributed across the sequence (Figure 3A, Supplementary Figure 8). These data were used to assess the ensemble of generated structures by quantifying the number of satisfied cross-links. In these analyses, distance constraints of 40 Å between the Cα of two lysines were considered, imposed by the spacer arm length of the DSBSO cross-linker, while accounting for some degrees of domain and side-chain flexibility (Zhu et al., 2024). Notably, 57% of the observed cross-links were compatible with the initial model before the MD simulations, with the alignment rising to 74% when the flexible IDR was excluded from the analysis (Figure 3B, C, Supplementary Figure 9A). Compatible cross-links were primarily observed within the domains, while intra-domain cross-links remained out of range due to the extended domain-linker conformations (Figure 3C, yellow for in range, red for out of range). Multiple valid inter-domain cross-links were detected between residues brought in proximity by domain homo-oligomerization, thereby confirming the assemblies observed in the crystal structures (Supplementary Figure 9B). After the MD simulations, the structural model and the experimentally derived cross-links agreed further, as nearly all identified cross-links (97%) were satisfied at some point during the MD simulations (Figure 3B). Notably, in both scenarios, meaning whether all cross-links were considered or those involving the IDR were excluded, MLCG simulations showed improved agreement with the experimental data at shorter simulation times compared to all-atom simulations (Figure 3B, Supplementary Figure 6C).

**Figure 3:**
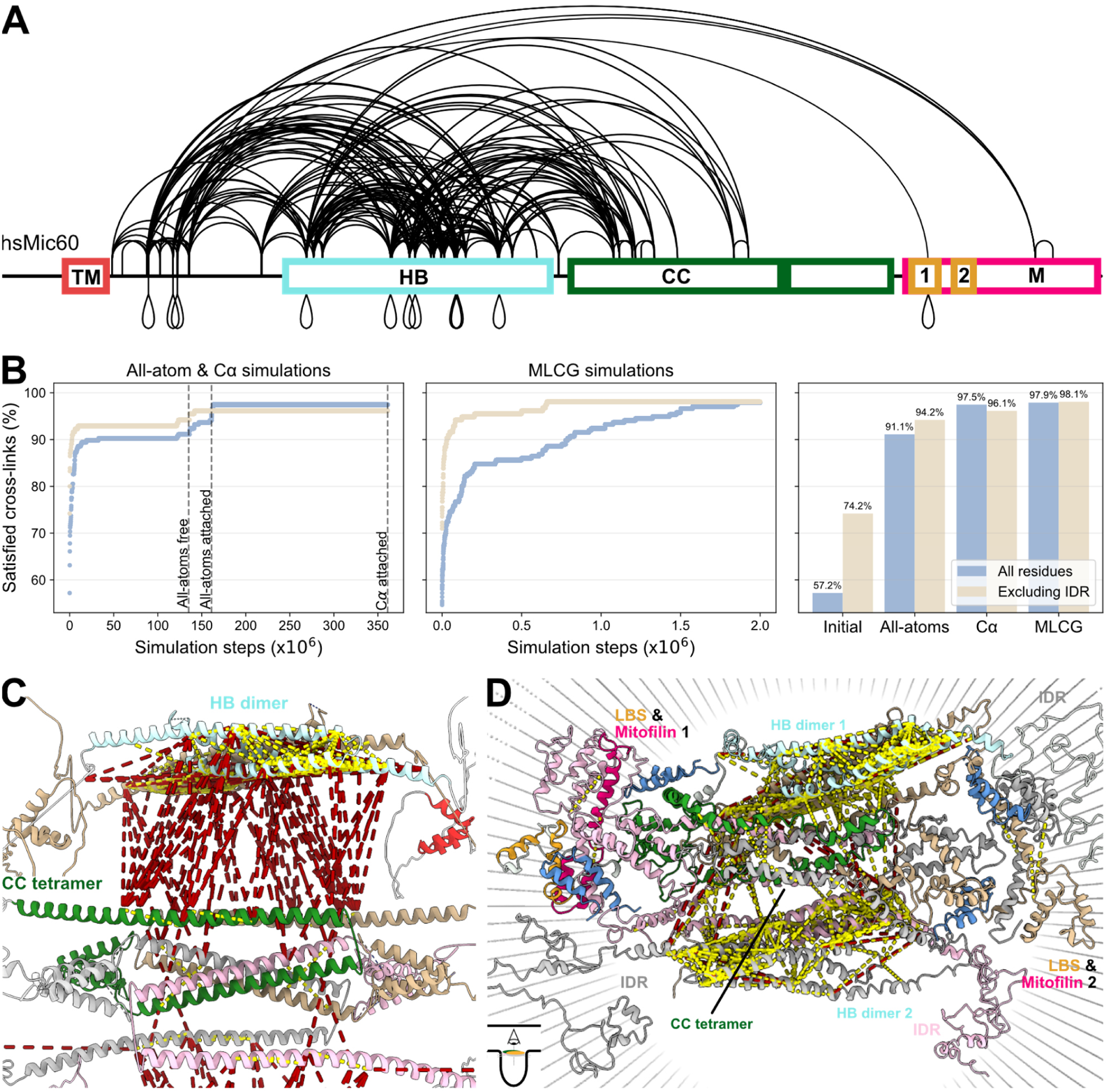
*In organello* cross-linking as model validation. **A** Visualisation of the DSBSO mitochondrial cross-linking dataset on the Mic60 sequence, modified from xiNET (Combe et al., 2015). Cross-links between different lysines are indicated above whereas self cross-links are indicated below the domain scheme. **B** The percentage of cross-links satisfied across the frames of the simulation, shown for the all-atom and Cα pipeline (left) and the MLCG model (center). The MD simulations were separated into the initial equilibration of the all-atom simulation prior to attachment to the crista (“all-atoms free”, first 135 million steps), the equilibration of the all-atom model attached to crista (“all-atoms attached”, 25 million steps), and the Cα-based simulations attached to the crista (“Cα attached”, 200 million steps). Many of the cross-links involve the flexible IDR and were therefore excluded and the dataset was reanalysed (“Excluding IDR”). The rightmost panel shows the percentage of cross-links from the published DSBSO dataset observed in the human Mic60-Mic19 model before (“Initial”), after the all-atoms simulations (“all-atoms”), after the subsequent Cα simulations (“Cα”), and using the MLCG model. Satisfied cross-links were computed considering all parallel simulations cumulatively across the duration of the simulations (n=30 for all-atom and Cα and n=6 for MLCG simulations). **C, D** Closeup view of the homology-based model of the human Mic60-Mic19 complex shown before (C, as in Figure 2A) and after (D, as in Figure 2D) the MD simulations. The cross-links annotated in A are visualized as dashed lines connecting the corresponding lysine residues (yellow for in range, red for out of range, cut-off set to 40 Å). All cross-links were visualized on the Mic60-Mic19 model using ChimeraX with the XMAS tool (Lagerwaard et al., 2022; Meng et al., 2023).

The overall increase in agreement after the MD simulations compared to the initial model was attributed on one hand to the collapse of the HB on the CC core during the all-atom simulations, rendering multiple intra-domain cross-links valid. On the other hand, the Cα-based simulations reflecting the dynamic conformations of the IDR further improved the fit of the model to the cross-linking data (Figure 3B, D). Thus, the *in organello* cross-linking results further support the notion of a highly dynamic model, in which the dimeric HB tightly packs against the Mic60 CC tetrameric core, with the flexible IDR surrounding them. The cross-linking validation of HB-CC packing favours the model presented here over the alternative model, which showed markedly lower satisfied cross-links during all-atom simulations (Supplementary Figure 5E, Supplementary Figure 10).

### The Mic60 subcomplex as a diffusion barrier in human mitochondria

CJs were suggested to function as a diffusion barrier that regulates the exchange of proteins and metabolites between the cristae lumen and the IMS (Gottschalk et al., 2022; Rampelt et al., 2017). While the Mic60 subcomplex is the largest assembly with a soluble moiety in the CJ pore, direct experimental evidence supporting a role as a diffusion barrier has remained elusive. To investigate this idea computationally, we leveraged our dynamic model of the human Mic60-Mic19 complex to probe its capacity to restrict molecular diffusion. In these simulations, we assessed the ability of spherical particles of varying radii to traverse the CJ plane from the IMS into the cristae lumen (Figure 4A, Supplementary Video 1). The MD simulations revealed a steady decrease in the flux of particles crossing the Mic60-Mic19-filled protein pore with increasing particle size. Crossing was efficiently excluded for particles with a radius larger than 2 nm (corresponding to a globular protein of ∼27 kDa (Fischer et al., 2004)). With smaller particle sizes, crossing was gradually observed (Figure 4B).

**Figure 4:**
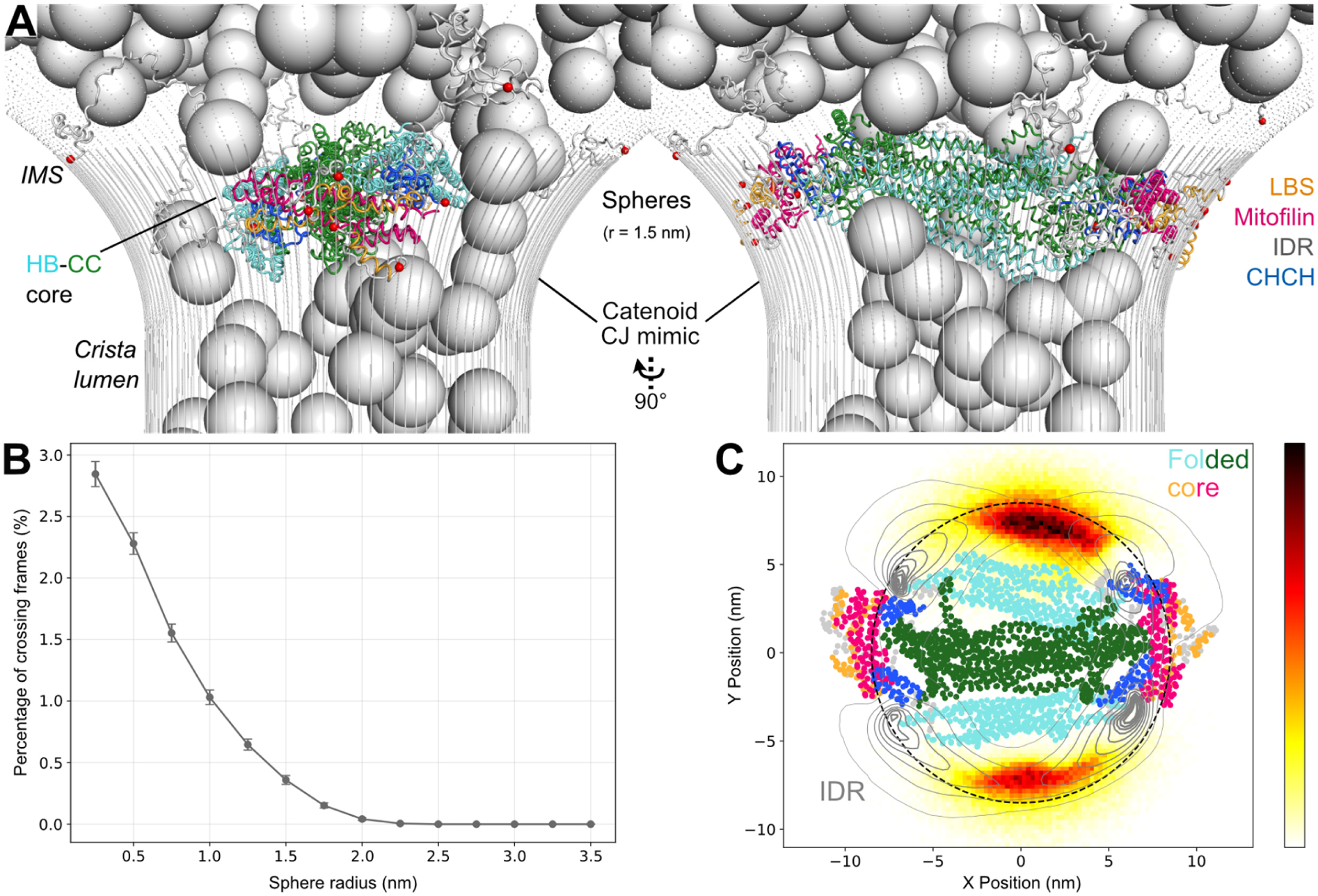
The human Mic60-Mic19 complex as a diffusion barrier in CJs. **A** Snapshot of the Cα-based diffusion experiments of the human Mic60-Mic19 model in the catenoid. 200 spheres, here with the radius of 1.5 nm, were allowed to pass through the junction. There were no repulsion forces between the spheres, only between the spheres and the complex or the catenoid, allowing them to be simulated simultaneously. **B** Fraction of frames exhibiting an above-to-below Mic60-Mic19 plane transition at equilibrium, normalized by the total number of frames, plotted as a function of sphere radius. Grey bars indicate the mean and standard deviation across all spheres. **C** Heat map depicting the density of sphere trajectories (radius 1.5 nm) projected onto the xy-plane during crossing events through the CJ in the presence of the human Mic60-Mic19 complex. Regions in black indicate the highest crossing density, while white indicates the lowest. The position of the complex throughout each simulation frame is projected onto the CJ plane. The average structure of the folded domains is displayed as a cartoon, while the IDR is illustrated as black contour lines to better illustrate its movement. The dashed black line indicates the CJ diameter of 17 nm.

We previously suggested that the IDR of Mic60, a common feature across Mic60 orthologues, may play a role in impeding the passage of larger molecules (Bock-Bierbaum et al., 2022). To test this hypothesis, we analysed a heat map of crossing events for particles at a radius of 1.5 nm, stratifying the complex into two functional regions: the folded core (displayed as a coloured cartoon) and the IDR (grey contour lines) (Figure 4C). Projection of these regions across all simulation frames demonstrated that the dynamic movement of the IDR enables it to intermittently occupy wider sections of the catenoid-shaped membrane, thereby extending its spatial coverage beyond that of the structured core. As a result, the IDR effectively limited access through the CJ plane. When passing the IDR, particle passage was confined to the narrow strait between the folded complex core and the CJ membrane, severely restricting diffusion. Our simulations therefore suggest that the dynamic nature of the Mic60-Mic19 subcomplex and particularly the dynamics of the IDRs facilitate its function as a diffusion barrier.

## Discussion

Our study provides structural and mechanistic insight into the highly dynamic architecture of the human Mic60-Mic19 subcomplex through an integrative structural biology approach combining *in vitro*, *in silico* and *in organello* data. Our MD simulations further indicate that the dynamic nature of the Mic60-Mic19 subcomplex is an important feature for its function as a mitochondrial diffusion barrier.

We identified the Mic60 HB as a previously uncharacterized oligomerization domain specific to animal orthologues, thus completing the structural annotation of the folded domains in human Mic60. Notably, the Mic60 HB forms an antiparallel dimer composed of three-helix bundles, reminiscent of the architecture adopted by the apoptotic signalling protein Smac/DIABLO (Chai et al., 2000; van Kempen et al., 2024). Smac/DIABLO localizes to the IMS in non-apoptotic conditions and was identified as a MICOS interactor in the MINDNet network (Adrain et al., 2001; Schaumkessel et al., 2025), suggesting that this fold may represent a conserved structural domain in IMS proteins and function as an interaction platform.

A direct physiological role of the Mic60 HB in establishing CJ architecture is supported by observations in cardiomyocytes, which express a Mic60 splice variant lacking exon 6, corresponding to a deletion of residues 176-208, which include the N-terminal part of the HB (Zhu et al., 2021). While the missing residues do not directly contribute to HB dimerization based on our structure, the deletion could disrupt interactions with adjacent IDR residues or promote local unfolding and could therefore be linked to mitochondrial phenotypes observed in these cells, including elongated morphology and densely packed cristae (Li et al., 2020; Piquereau et al., 2013).

To fully leverage the available structural and biochemical information, we manually constructed an initial model of the human Mic60-Mic19 complex using experimentally determined crystal structures as basis. Each domain was assumed and modelled to form homo-oligomers consistent with crystallographic and biochemical data: a HB dimer, a CC tetramer, and a mitofilin dimer. While both HB dimerization and CC tetramerization were experimentally addressed for the human orthologue, expression of the isolated human mitofilin domain was not successful, precluding similar biochemical validation. However, *in organello* cross-links from purified human mitochondria supported the proposed homo-oligomerization of all three domains, in agreement with the corresponding crystal structures of fungal homologues. We further assumed that the human Mic60-Mic19 complex spans across the CJ neck, rather than spanning from the IMM to the OMM. A published computational model proposed that Mic60 functions as a monomer, anchored to the IMM via its TM domain and extending across the IMS to interact with the flat OMM with its mitofilin domain (Brown et al., 2025). Contrasting this idea, the symmetric architecture required by Mic60’s antiparallel tetrameric state and the inclusion of Mic60’s TM domain in the IMS are not compatible with a monomeric, asymmetric IMM-OMM bridging role. Furthermore, our *in silico* analysis indicates that the mitofilin dimer exhibits strong binding to a curved membrane surface mimicking the IMM. In line with the model suggested for the fungal MICOS orthologues (Bock-Bierbaum et al., 2022), we therefore propose that the Mic60-Mic19 complex adopts a CJ-spanning architecture, with the tetrameric HB-CC core occupying the CJ pore and the mitofilin domains contacting the IMM via their lipid binding sites. The HB dimer, in the context of the fully assembled complex, significantly increases the CJ interface occupied by the folded complex by packing against the tetrameric CC, thereby also contributing to restricting diffusion in the tubular CJs of animalia.

According to our integrative model, the Mic60-Mic19 hetero-octamer spans across the CJ at a diameter of 17 nm, with the flexible IDRs of Mic60 moving extensively in the CJ opening. Our diffusion simulations suggest the exclusion of proteins with a hydrodynamic radius of ∼2 nm (∼27 kDa). This would exclude larger assemblies like small TIM heterooligomers from crossing the CJ, and possibly also smaller globular proteins such as cytochrome c (PDB ID: 1J3S) which are confined to the cristae (Mühlenbein et al., 2004; Scorrano et al., 2002). It can be envisaged that the inclusion of further MICOS components into our model, such as the coiled-coil domains of Mic19 or the membrane-embedded Mic10 subcomplex, will further restrict the size of particles that can cross the CJ pore. We furthermore show that the flexibility of the IDR is a prerequisite for its function as a diffusion barrier through the crista junction. In our current simulations, the interaction of the IDR with the particles was repulsive. In contrast, the IDR may contribute to cargo selectivity via low affinity interactions with the substrates. In an analogous way, phenylalanine-glycine repeats of the nuclear pore complex have been shown to bind to the human immunodeficiency virus allowing its efficient translocation into the nucleus (Dickson et al., 2024).

The dynamic MICOS model introduced in this study could be integrated into larger-scale simulations of IMM protein organization, a challenge noted in previous efforts to simulate MICOS due to its inherent complexity and flexibility (Brown et al., 2025). The applied integrative approach may provide a valuable benchmarking strategy for modelling proteins with IDRs, which pose a persistent challenge in structural biology due to their conformational heterogeneity. IDRs frequently mediate transient interactions and contribute to structural flexibility – features especially relevant in dynamic, membrane-associated assemblies that regulate diffusion, such as the nuclear pore (Winogradoff et al., 2022). Our approach builds on these insights by validating such simulations with *in organello* data, a concern often raised about *in silico* measurements (Rush et al., 2023). By integrating experimental constraints with flexible modelling, we offer a framework for capturing the structural and functional contributions of disordered regions within large macromolecular complexes, while physiological validation anchors the model in a relevant cellular context. Our integrative human Mic60-Mic19 model therefore sheds light on the dynamic role of MICOS as a governor of CJ dynamics through a pipeline combining structural, biochemical, computational, and cross-linking data.

## Materials and Methods

### Cloning and plasmids

hsMic60 (UniProt ID: Q16891, cDNA synthesized by Eurofins Genomics) was used for cloning. The hsMic60 helical bundle (Mic60 HB, residues 198-378) was cloned via restriction sites *Nde*I and *Xho*I and subsequent ligation into pET SUMO vector encoding an N-terminal His_6_-SUMO tag fusion protein. Mutations were introduced via site-directed mutagenesis (Ho et al., 1989). The hsMic60 coiled-coil (hsMic60 CC, residues 473-542 and residues 473-617) was cloned into a pET21b(+) vector encoding a human rhinovirus 3C (HRV-3C) cleavable N-terminal MBP fused to a pMal_c2X linker using Gibson assembly.

### Expression and purification

Expression plasmids were transformed into BL21(DE3) *Escherichia coli* cells and plated on ampicillin-containing LB plates. Transformed bacteria were grown in terrific broth media supplemented with 100 μg/ml ampicillin at 37 °C and 80 rpm until they reached an OD_600_ of 0.8 and subsequently cooled to 18 °C and protein expression was induced with 300 μM isopropyl-ß-D-1-thiogalactopyranoside (IPTG) overnight. Cells were harvested via centrifugation for 20 min at 4,000 g and cell pellets were stored at −20 °C.

Mic60 HB pellets were thawed and resuspended in lysis buffer (20 mM 4-(2-hydroxyethyl)-1-piperazineethanesulfonic acid (HEPES/NaOH pH 7.5, 500 mM NaCl, 20 mM imidazole), supplemented with 500 μM 4-(2-aminoethyl)-benzolsulfonyfluorid hydrochloride (AEBSF, BioChemica) and 1 μM DNase (Roche). Cells were lysed through sonication (5x 45 sec with 1 sec pulses at 70% power) and lysates were centrifuged at 40,000 g for 45 min. The supernatant was filtered using a 0.45 μM filter and applied to Ni-NTA resin, equilibrated in lysis buffer. The column was washed with 5 column volumes (CV) lysis buffer and 5 CV wash buffer (20 mM HEPES/NaOH pH 7.5, 500 mM NaCl, 40 mM imidazole) and the protein was eluted in 5 CV elution buffer (20 mM HEPES/NaOH pH 7.5, 500 mM NaCl, 500 mM imidazole). His_6_-tagged SUMO Protease was added at a 1:50 ratio (w/w) and the protein mixtures were dialyzed overnight at 4 °C to low imidazole buffer (20 mM HEPES/NaOH pH 7.5, 150 mM NaCl, 20 mM imidazole). The protein was subsequently applied to a reequilibrated Ni-NTA column to remove the residual His_6_-SUMO tag and SUMO protease. The flow-through was collected, concentrated using an Amicon filter concentrator of 10 kDa cut-off (Millipore) and applied on a HiLoad™ 16/600 Superdex™ 75 preparative grade column connected to an ÄKTA Pure chromatography system (Cytiva). Protein was eluted at a flow-rate of 1 ml/min and 1 ml fractions were collected. Pure protein fractions were pooled, concentrated, flash-frozen in liquid nitrogen and stored at −70 °C. Point mutants were purified using the same protocol, however for artificial cysteine variants IMAC column buffers were complemented with 2 mM β-mercaptoethanol and the SEC buffer was complemented with 2 mM dithiothreitol (DTT) to hinder unspecific disulfide bridge formation.

Mic60 CC bacterial pellets were resuspended in lysis buffer (20 mM HEPES/NaOH pH 7.5, 500 mM NaCl, 1 mM Ethylenediaminetetraacetic acid (EDTA)), supplemented with 500 μM AEBSF and 1 μM DNase, lysed through sonication and centrifuged as described for Mic60 HB. The filtered supernatant was applied on Dextrin Sepharose HP beads (Cytiva), washed with 5 CV of lysis buffer and eluted using lysis buffer supplemented with 10 mM maltose. The eluate was concentrated using an Amicon filter concentrator of 30 kDa cut-off and applied on a Superose™ 6 Increase 10/300 GL column connected to an ÄKTA Pure chromatography system (Cytiva). The protein construct CC_1 (hsMic60 residues 473-542) eluted in an earlier shoulder peak at volume 14.2 ml and a main peak at volume 15.3 ml. The two peaks were concentrated separately as samples “shoulder” (3 mg/ml) and “main” (20 mg/ml). For the construct hsMic60 CC_2 (hsMic60 residues 473-617), only one peak was observed and concentrated to 35 mg/ml.

For all protein samples, purity was confirmed by mass spectrometry as described previously (Bock-Bierbaum et al., 2022) and samples were flash-frozen in liquid nitrogen and stored at −70 °C.

### Crystallization, data collection, refinement and structural analysis tools

Mic60 HB crystallization trials were performed using the sitting-drop vapor-diffusion method in a 96-well crystallization plate using a Gryphon pipetting robot (Matrix Technologies Co.) by mixing 200 nl protein with 200 nl reservoir solution and equilibrating against 80 μl reservoir. Commercial screens, namely JBScreen Basic HTS, JBScreen JCSG+ HTS, JBScreen PEG/Salt (all Jena Bioscience), Classics II Suite, pH Clear Suite I and II, Protein Complex Suite (Qiagen), were used for initial screening at 4 °C and 20 °C using a concentration of 13 mg/ml. The Rock Imager 1000 storage system (Formulatrix) was used for storing and imaging the experiments. Initial hits were acquired for Mic60 HB (19-23 mg/ml) at 25% w/v PEG 3350, 0.1 M Tris/HCl pH 8.5 after incubation at 4 °C for one day. Optimization of PEG concentration (15.7-19.4% w/v) and pH (8.3-8.8) yielded protein crystals up to 300 μM in length, but with poor diffraction to 6-8 Å. Low diffracting crystals were used as seeds by mixing them with 120 μl 21.4% PEG 3350, 0.1M Tris/HCl pH 8.9 and 30 μl Mic60 HB (19 mg/ml) and crushing by vortexing using a microseeding bead for 1 minute (Shaw Stewart et al., 2011). Microseeding crystal trials against commercially available screens were set up at 4 °C using 200 nl Mic60 HB (19 mg/ml), 300 nl reservoir solution and 100 nl of a 1:10 dilution of the seeding solution in the sitting drop, equilibrated against 80 μl of reservoir solution. After 14 days, a crystal grown in 0.2 M ammonium citrate, 20% PEG 3350 was fished and flash-frozen in liquid nitrogen in presence of 15% ethylene glycol as cryoprotectant.

Diffraction data were collected at a wavelength of 0.9184 Å and −173 °C on beamline 14.1 operated by the Helmholtz-Zentrum Berlin at the BESSY II synchrotron in Berlin Adlershof (Mueller et al., 2025; Mueller et al., 2015). 1,600 images were collected using 0.25 s exposure time, 426.6 mm detector distance and an oscillation increment of 0.1°. Data was automatically indexed, integrated and scaled using XDSAPP V. 3.1.9c (Sparta et al., 2016). Mic60 HB crystallized in space group P4_1_2_1_2 (92). The crystallographic phase problem was solved by molecular replacement with Phaser-MR implemented in the Phenix distribution (V. 1.20.1_4487) using two copies of AlphaFold2 predicted hsMic60 residues 198-378 (Jumper et al., 2021; Liebschner et al., 2019; McCoy et al., 2007). The Mic60 HB structure was finally solved employing Phenix.autobuild (Terwilliger et al., 2008), followed by manual model building in Coot V. 1.1.11 (Emsley et al., 2010) and iterative model refinement with phenix.refine using non-crystallographic symmetry and Translation-Liberation-Screw rotation (Afonine et al., 2012). The final model contains two copies of Mic60 HB in the asymmetric unit, assembled in a dimer with non-crystallographic symmetry, with residues 210-261 and 271-377 visible for monomer 1 and 198-261 and 271-377 visible for monomer 2. Molprobity V. 4.0.2 (Williams et al., 2018) was used for final model validation, the according data statistic can be seen in Supplementary Table 1. The structure was deposited and validated in the PDB database under PDB ID 9QWR.

Figures of the Mic60 HB and of the Mic60-Mic19 model were generated using PyMol (V. 2.5.5) (Schrödinger & DeLano 2020). PDBePISA was used for interface analysis of the Mic60 HB dimer (Krissinel & Henrick, 2007). The FoldSeek server was employed to perform protein structure alignments of the Mic60 HB, using the construct amino acid sequence as input (van Kempen et al., 2024). For evolutionary conservation analysis of the Mic60 HB, Mic60 orthologues postulated to possess a HB were initially identified based on their AlphaFold2 predictions. From these orthologues, eight representative organisms for the kingdom of animalia were identified (UniProt accession number in brackets: *Homo sapiens* (Q16891), *Mus musculus* (Q8CAQ8), *Danio rerio* (Q6PFS4), *Xenopus laevis* (A0A1L8HKP3), *Apis mellifera* (A0A7M7GMI1), *Drosophila melanogaster* (P91928), *Caenorhabditis elegans* (IMMT-1 Q22505, IMMT-2 Q9XXN2), *Hydra vulgaris* (A0A8B7DIZ7)) and their amino acid sequences were aligned using the ConSurf Web server using default parameters (Yariv et al., 2023). It was manually confirmed that the sequence segments aligned to the Mic60 HB were the ones predicted by AlphaFold2 to form the HB of each orthologue. The aligned HB sequence segments were extracted and imported to Jalview (Waterhouse et al., 2009) to generate a conservation score plot and a phylogenetic tree for the Mic60 HB after Huynen et al. (2016). The ConSurf Server was then used, with the structure of the Mic60 HB dimer and the Mic60 HB sequence alignment as input, to generate structure conservation plots (Yariv et al., 2023).

### Oxidation of cysteines and non-reducing SDS electrophoresis

For the generation of fully reduced protein samples for disulfide bond analysis, the sample buffer was exchanged and DTT concentration was increased to 10 mM (20 mM Tris/HCl pH 7.5, 150 mM NaCl, 10 mM DTT). For the oxidation reaction, protein samples were dialyzed overnight to CuSO_4_ oxidation buffer (20 mM Tris/HCl pH 7.5, 150 mM NaCl). 0.1 mM CuSO_4_ was added to the samples (5 mg/ml) and then incubated on ice for 1 min. The reaction was quenched by addition of 50 mM EDTA and residual CuSO_4_ and EDTA were removed using a PD-10 column (Cytiva). 10 μl sample (0.5 mg/ml) was added to 2.5 μl non-reducing SDS loading buffer (250 mM Tris pH 6.8, 10% SDS, 30% glycerol). Gels were stained using Coomassie Brilliant Blue and imaged using a Gel Doc™ XR+ Gel Documentation System. Protein bands were quantified by pixel count using ImageJ (V. 1.53k) (Schneider et al., 2012). For statistical analysis out of three replicates t-test was used, assuming normal distribution, and standard deviation was calculated.

### Analytical size-exclusion chromatography coupled to right angle light scattering (SEC-RALS)

Analytical SEC-RALS was performed using an Agilent HPLC system equipped with an autosampler G1329B and an Omnisec Reveal (Malvern). A Superdex 75 Increase 5/150 GL column and a Superose 6 Increase 5/150 GL were used for Mic60 HB and Mic60 CC measurements accordingly. The column and RALS instrument were equilibrated overnight with SEC buffer (20 mM HEPES/NaOH pH 7.5, 150 mM NaCl). 100 μl sample (Mic60 HB 1 mg/ml, Mic60 CC 2 mg/ml) was injected using the autosampler and eluted at a flow rate of 0.2 ml/min at 20 °C. UV absorption at 280 nm, 260 nm, 210 nm and light scattering were measured and analysed by the Agilent Offline and Malvern Omnisec software.

### Blue Native polyacrylamide gel electrophoresis (BN-PAGE)

Recombinant proteins were analysed by BN-PAGE as described previously (Bock-Bierbaum et al., 2022). Briefly, 10 μg of purified protein were applied on a 4-16% acrylamide gradient BN-PAGE gel which was run at 150 V for 2 h on ice and subsequently stained using Coomassie Brilliant Blue.

### Assembly of the initial Mic60-Mic19 model

A model of the Mic60-Mic19 complex was assembled in order to study it using molecular dynamics simulations. Three solved structures were used as the basis for this model: the Mic60 HB (residues 198-378, described in this paper), the CC domain of *Lachancea thermotolerans* Mic60 (PDB ID: 7PUZ, residues 207-382, equivalent residues in hsMic60 are residues 443-603) and a fusion construct of the Mic60 mitofilin domain with the Mic19 CHCH domain from the organism *Chaetomium thermophilum* (PDB ID: 7PV1, ctMic60 565-586-GS-622-691 and ctMic19 residues 116-164, equivalent residues are hsMic60 residues 627-751 and hsMic19 residues 180-222).

To utilize the previously published fungal orthologue structures in assembling the human Mic60 model, homologous structures were generated using SWISS-MODEL, covering the residue range included in the experimental models (Waterhouse et al., 2018). According to AlphaFold2 (Jumper et al., 2021), the remaining sequence segments were predicted to be the N-terminal intrinsically disordered region (residues 1-197, including the transmembrane helix), an extended helical connection between the Mic60 HB and the CC (residues 379-442), short unfolded linker regions and the flexible LBS2 (residues 649-685). The majority of these segments, apart from the helical connection, was predicted to be unstructured and included in the final model according to the AlphaFold2 prediction of full-length hsMic60.

Structural information for hsMic19 is only available for the CHCH domain, leading us to exclude the remaining protein sequence in the final Mic60-Mic19 model to omit inaccurate model building. The SWISS-MODEL generated homologue was modelled on the mitofilin domain of the hsMic60 model according to the previously published structure on their interaction (PDB ID: 7PV1).

The SWISS-MODEL orthologues and AlphaFold2 predictions were manually integrated using Coot and spatially adjusted using iterative cycles of ChimeraX V. 1.9 (Meng et al., 2023), ultimately assembling a model of the human Mic60-Mic19 hetero-octamer. During this process, particular attention was paid that the structural elements retain their experimentally determined characteristics, e.g. their oligomerization state. Some short structural segments between the CC and the LBS1 (residues 527-536, 555-561, 584-591 and 603-607) predicted by AlphaFold2 to be helical were reconstructed as flexible linkers in the molecular dynamic simulations.

While the model reflects the known structural information on individual domains of hsMic60, there is no experimental data on the relative positioning of these domains to one another. The oligomeric state of the individual domains results in the CC forming a tetrameric core, as illustrated previously for fungal Mic60 (Bock-Bierbaum et al., 2022). The short linker length to the LBS-mitofilin domain as well as the directionality of the LBS domains pointing away from the complex and towards the hypothetical inner membrane lead to a planar orientation for these domains of the tetramer. Neither experimentally nor computationally could it be determined how the hsMic60 CC tetramer is connected to the hsMic60 HB dimers and how the HB dimers are positioned relative to the tetrameric CC. Therefore, two models were generated. In model 1, the HB domains and mitofilin domains dimerized via different Mic60 molecules, whereas in model 2, the same molecules mediate dimerization of the two domains (Supplementary Figure 4C, D).

### Molecular dynamics simulations of the Mic60-Mic19 model

For all the simulations, the N-terminal region of Mic60 exposed to the matrix as well as the transmembrane domain, corresponding to the first 61 residues, was removed, making Ile62 the N-terminus of cristae-soluble Mic60. To fill the sequence gaps and the overstretched regions, a homology modelling process was performed using Modeller (Webb & Sali, 2016) and a 4-fold structural symmetry was subsequently applied to the initial model, using a custom protocol to apply symmetry restraints to the Cα atoms. The Modeller MD refinement protocol was performed at the *slow* level. A total of 300 models were generated, picking the one with the lowest DOPE score (Shen & Sali, 2006).

All-atoms molecular dynamics simulations of the Mic60-Mic19 complex were set up using OpenMM (Eastman et al., 2024). The system was aligned along its principal axis and immersed in solvent within a simulation box measuring 27.5 × 19.8 × 15.3 nm^3^. The solvent consists of TIP3P water molecules neutralized with Na^+^ and Cl^−^ ions at 0.15 M ionic strength (Jorgensen et al., 1983). Non-bonded interactions are treated with Particle Mesh Ewald, cut-off of 1 nm and a 0.75 nm switching distance. Hydrogen bonds are constrained, and hydrogen mass was set to 4 atomic mass units (amu) for stability. The force filed employed is the Amber14SB force field (Maier et al., 2015). The system was slowly equilibrated using a multi-step protocol to ensure stability under physiological conditions. The system was subjected to energy minimization before gradual thermal and pressure equilibration with Langevin Middle Integrator and a 2 fs timestep. Harmonic positional restraints of 500 kJ/mol/nm² were initially applied to the protein heavy atoms. Equilibration began with an NVT warm-up phase (NVT for number (N), volume (V), and temperature (T)) lasting 200 ps, during which the temperature was gradually increased from 10 K to 300 K over 100,000 steps. The system then underwent NPT equilibration for 400 ps with a Monte Carlo barostat set at 1 bar and 300 K, maintaining the initial restraints throughout 200,000 steps. To progressively relax constraints, the harmonic restraint force was then reduced from 500 kJ/mol/nm² to 100 kJ/mol/nm² over 400 ps gradually. Positional restraints were removed from side chains and applied exclusively to the backbone, maintaining a force constant of 100 kJ/mol/nm² for 400 ps. Finally, the backbone restraints were gradually reduced to zero over 600 ps. Positional restraints were removed from side chains and applied exclusively to the backbone, maintaining a force constant of 100 kJ/mol/nm² for 400 ps. Finally, the backbone restraints were gradually reduced to zero over 600 ps.

Starting from the equilibrated structure, 30 simulations were run to maximize the exploration of the conformational space, increasing the timestep to 4 fs and saving frames every 0.04 ns. Simulations were run for around 500 ns until RMSD equilibration with respect to the common initial structure, as shown in Figure 2C.

For the alternative model, the same procedure described above was applied. The system was solvated in a 36.6 × 27.6 × 20.2 nm simulation box, larger than that used for the main model, to accommodate its increased longitudinal dimension. Three independent simulations were subsequently performed, each extending for approximately 170 ns.

Using OpenMM’s CustomExternalForce, each of the 30 final structures was gradually coupled to an external potential that emulates a crista junction, modelled as a concatenation of a cylindrical and catenoid geometry (Eastman et al., 2024). The CJ was defined with a diameter of 17 nm. Attachment of Mic60 was achieved at the N-terminus by linking the Cα atom of the first residue (I62) to the potential. For the C-terminus, two attachment points per chain were selected, corresponding to the Cα atoms of residues flanking the LBS2 region (Arg652, Pro670). These points were tethered via a harmonic potential with a force constant of 200 kJ/mol/nm² applied perpendicular to the crista surface, allowing free tangential mobility along the crista. The coupling was implemented incrementally, with the force constant ramped from 0.1 kJ/mol/nm² to 200 kJ/mol/nm² over 10 steps of 5 ns each. Following the equilibration phase of the attachments, a production simulation of 100 ns was conducted.

MLCG simulations were performed using the chemically transferable model published in (Charron et al., 2025). This model employs a coarse-grained representation with a resolution of five beads per residue. All information required to run simulations with the *mlcg* package is publicly available in the Code Availability section of (Charron et al., 2025). For the Mic60–Mic19 complex, four independent simulations were conducted, each consisting of 2,000,000 timesteps. Simulations were carried out using a Langevin integrator at a temperature of 300 K, with a friction coefficient of 1.0 and a timestep of 4 fs. Positions were saved every 10 timesteps.

### *In silico* force calculation of Mic60-Mic19 binding to the IMM

To evaluate the interaction strength of the complex with the IMM, we conducted coarse-grained molecular dynamics simulations using the Martini force field (Souza et al., 2021) and performed umbrella sampling to quantify the free energy of binding. We focused on the Mic60-Mic19 C-terminal mitofilin-CHCH lobe, which was prepared using the martinize2 script with the Martini 3 force field, incorporating a Go-like network to preserve secondary structure and native contacts (Souza et al., 2024). The system was solvated using the insane script (Wassenaar et al., 2015), with a simulation box size of 17 × 23 × 35 nm³ and a lipid composition of POPC:POPE:CDL0:POPS in a 40:40:15:5 ratio to mimic the IMM (Basu Ball et al., 2018). Na⁺ and Cl⁻ ions were added to achieve an ionic concentration of 0.15 M. The system was then simulated for 1 µs using martini_openmm (MacCallum et al., 2023) at 300 K, with a friction coefficient of 10 ps⁻¹, a timestep of 20 fs, a nonbonded interaction cut-off of 1.1 nm, and a barostat set to 1 bar. To simulate the curvature of the crista membrane, we first removed the solvent and applied membrane bending using the expression described in (Mahmood et al., 2019), with a γ factor of 0.05. The system was then re-solvated to accommodate the new membrane dimensions. To preserve the membrane curvature, the barostat was set to 1 bar, acting exclusively along the z-dimension. Additionally, an OpenMM CustomExternalForce was implemented to apply a harmonic potential with a force constant of 200.0 kJ·mol⁻¹·nm⁻² on the headgroups of the lower leaflet. The system was subsequently equilibrated for 1000 ns using the same simulation parameters as described above.

The initial window frames for umbrella sampling were generated using steered molecular dynamics. The protein was pulled 10 nm away from the membrane using a CustomCentroidBondForce between the center of mass (COM) of the protein and the membrane, with a force constant of 1000.0 kJ·mol⁻¹·nm⁻² and a pulling speed of 1 × 10⁻⁵ nm/ps. To constrain lateral motion, a harmonic restraint was applied along the x and y directions to the COM of the protein, with a force constant of 200.0 kJ·mol⁻¹·nm⁻². Initial configurations along the reaction coordinate were extracted by defining umbrella sampling windows separated by 0.1 nm. Starting from the equilibrated position, five windows were placed in the direction of the protein closer to the membrane, while the remaining 50 were positioned progressively further away. Each window was then simulated for 1000 ns. The COM distance time series was analyzed using the weighted histogram analysis method (WHAM) (Grossfield, Alan, “WHAM: the weighted histogram analysis method”, V. 2.1.0, http://membrane.urmc.rochester.edu/wordpress/?page_id=126). As shown in Supplementary Figure 7A, the free energy difference upon binding is approximately 100 kJ/mol, which was used to parameterize the interaction strength of the attachment points in the crista potential described above.

### *In organello* cross-linking as validation for the Mic60-Mic19 model

Cross-linking mass spectrometry data from Zhu et al. (2024) (accessible through PRIDE accession number PXD046382) were employed to validate the ensemble of generated structures. Cross-link identifications derived from the standalone version of XlinkX (Liu et al., 2017) were filtered to an n_score of < 1e^−15^ and only intra-links within IMMT were retained if they were confirmed by at least 2 spectral evidences. The search engine output was converted into a ChimeraX XMAS compatible output (Lagerwaard et al., 2022) using R and then visualized on protein models. To speed-up conformational explorations, Cα structure-based simulation were started from the 30 all-atoms final structures (Clementi et al., 2000). The standard Cα structure-based angle potential was replaced with a restricted angle potential for stability (Bulacu et al., 2013). Native contacts were computed for each starting configuration using Shadow algorithm (Noel et al., 2012). Native contacts formed involving the IDR part for all the chains were excluded. Simulations were run for 100 ns with a timestep of 0.5 fs and a reduced temperature of 0.5 (Jackson et al., 2015). Cross-links were plotted on the hsMic60 sequence using xiNET (Combe et al., 2015).

The cross-link satisfaction of the alternative model was lower than that of the main model (Supplementary Figure 5E), leading us to prioritize the main model in subsequent analyses.

### *In silico* CJ diffusion experiments

The trajectory that satisfied the highest number of cross-linkers was selected for diffusion-like simulations. In the Cα structure-based system, 200 identical spheres, each with the same radius, were introduced above the Mic60-Mic19 plane. A harmonic potential was applied to confine the spheres within the crista junction potential. To impose additional constraints, upper and lower planes were set to restrict the spheres within a closed space. No interaction was set between the spheres, while a Lennard-Jones repulsion term, proportional to 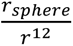, was introduced with Mic60-Mic19 and with the crista potential. The crossing event frequency was defined as the fraction of frames exhibiting an above-to-below Mic60-plane transition at equilibrium, normalized by the total number of frames, and was studied in correlation to the sphere radius. To approximately correlate particle radius to protein molecular weight we used the spherical approximation with a density of 1.35 g/cm^3^ (Fischer et al., 2004).

## Acknowledgements

We gratefully acknowledge funding from the Deutsche Forschungsgemeinschaft DFG (SFB/TRR 186, Project A12 (CC) and A23 (OD); SFB 1114, Projects B03, B08, and A04), the National Science Foundation (PHY-2019745), the Einstein Foundation Berlin (Project 042081510), the German Ministry for Education and Research (BMBF) project FAIME 01IS24076 and the MDC graduate school (EN). The authors gratefully acknowledge the computing time provided on the supercomputer Lise at NHR@ZIB as part of the NHR infrastructure and the BESSY team, Y. Roske for support during crystallographic data collection at beamline MX14.1, and T. Bock-Bierbaum, A. Natarajan, M. van der Laan and K. van der Malsburg for discussions. We are grateful to A. Schütz (Protein Production and Characterization Platform, MDC) for support with the mass spectrometry validation and H. Hashimi for his valuable insight into the HB conservation analysis.

## Contributions

EN designed constructs, solved the HB domain structure, performed the biochemical experiments and assembled the main and alternative Mic60-Mic19 models. ER set up and performed the MD simulations, with support from IZ. MR and EN contributed to the analysis of the cross-linking data along with ER. EN, ER, MR, IZ, FL, CC and OD designed research and interpreted structural, mass spectrometry and computational data. FL, CC and OD supervised research. EN and OD wrote the manuscript, with inputs from all authors.

## Additional information

### Author information

The atomic coordinates of Mic60 HB have been deposited in the Protein Data Bank with accession number 9QWR.

