## Supplementary Materials for "Integrative modelling reveals the structure of the human Mic60-Mic19 subcomplex and its role as a diffusion barrier in mitochondria"

### Supplementary Figures and Tables

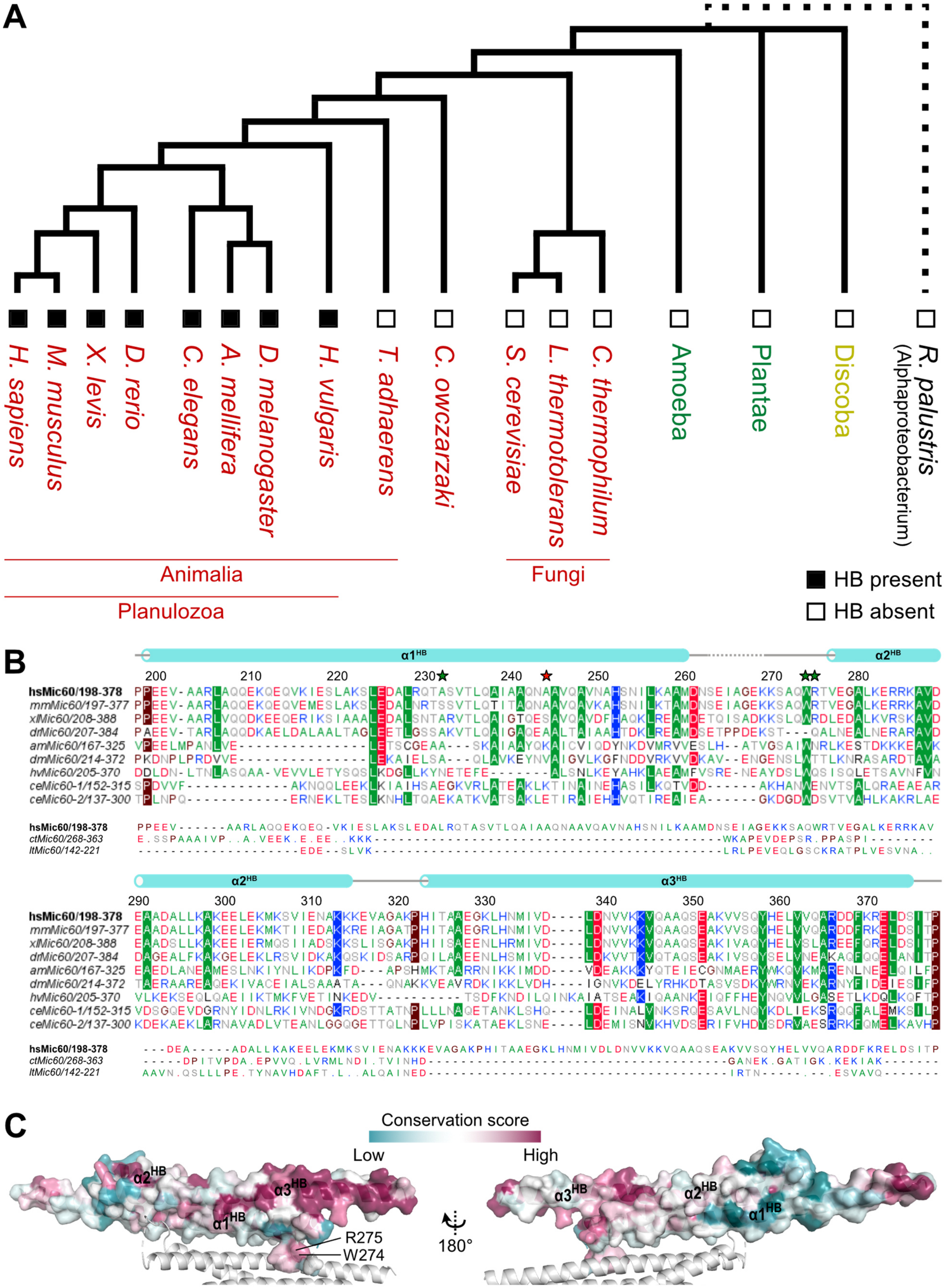

**Supplementary Figure 1: Evolutionary conservation analysis of the N-terminal helical bundle**

**A** Phylogenetic tree depicting the taxonomical relationship among the examined organisms (UniProt accession number in brackets): *Homo sapiens* (Q16891), *Mus musculus* (Q8CAQ8), *Xenopus laevis* (A0A1L8HKP3), *Danio rerio* (Q6PFS4), *Caenorhabditis elegans* (IMMT-1 Q22505, IMMT-2 Q9XXN2), *Apis mellifera* (A0A7M7GMI1), *Drosophila melanogaster* (P91928), *Hydra vulgaris* (A0A8B7DIZ7), *Trichoplax adhaerens* (B3S3B5), *Capsaspora owczarzaki* (A0A0D2UAX3), *Saccharomyces cerevisiae* (P36112), *Lachancea thermotolerans* (C5E325), *Chaetomium thermophilum* (G0SHY5). Dotted line shows unsure rooting of mitochondria with alphaproteobacteria. Organisms that possess the HB are marked with a black square, organisms expressing Mic60 but not predicted to possess a HB are marked with an empty square. **B** Sequence conservation analysis of the hsMic60 HB shows high sequence similarity among vertebrates (*Homo sapiens*, *Mus musculus*, *Danio rerio, Xenopus laevis*) and partial sequence preservation in protostomia (*Apis mellifera, Drosophila melanogaster*, *Caenorhabditis elegans*, *Hydra vulgaris*). A sequence alignment of hsMic60 with fungal orthologues (abbreviation and Uniprot accession number in brackets) illustrates they lack a HB and show low sequence similarity to the equivalent human sequence, shown here for *Chaetomium thermophilum* (ctMic60, G0SHY5) and *Lachancea thermotolerans* (ltMic60, C5E325). Highly conserved residues (> 60%, calculated based on orthologues where the HB is present) are placed in coloured boxes, individual residues are coloured according to their physicochemical properties (hydrophobic in green, positively charged in blue, negatively charge in red, prolines and glycines in brown, remaining in grey). Mutated residues are indicated by green stars, artificial cysteine bridge indicated by red star. **C** Surface representation of the Mic60 HB coloured by the ConSurf conservation score (cyan-to-purple colouring variable-to-conserved). The W274-containing loop and a patch of $\alpha3^{HB}$ comprise highly conserved surfaces in the HB, pointing to their functional relevance.

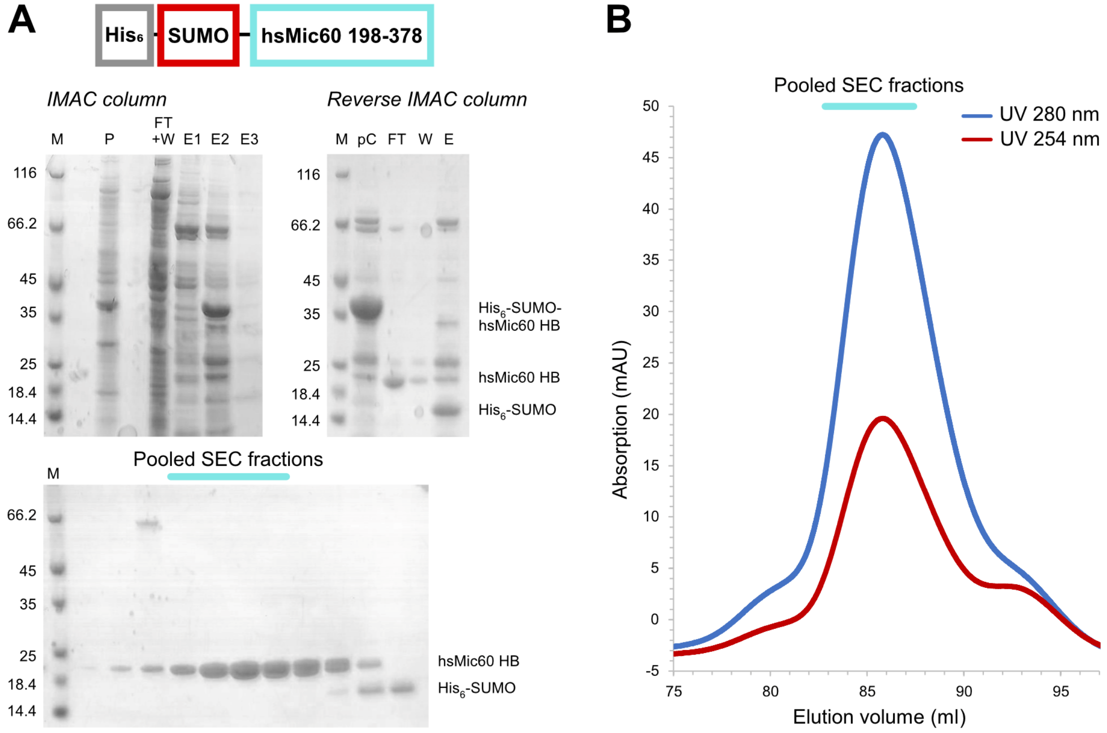


**Supplementary Figure 2: Purification of the Mic60 helical bundle (Mic60 HB)**

**A** Schematic overview of the Mic60 HB with an N-terminal $His_6$-SUMO tag. The protein was purified by Immobilized Metal Affinity Chromatography (IMAC, left SDS gel), followed by overnight cleavage using SUMO protease and a further IMAC (right SDS gel). Descriptions on the right side of the gel indicate the approximate molecular weight of the target protein and the $His_6$-SUMO tag. The final pure sample was acquired after SEC elution (bottom SDS gel). The cyan bar indicates the pooled fractions. P – Pellet, FT – Flow through, W – Wash, E – Eluate, pC – pre-cleavage. **B** Chromatogram of the SEC elution profile of Mic60 HB. Absorption at 280 nm (dark blue) and 254 nm (red) are plotted over the peak elution volume. The cyan bar indicates the pooled fractions shown in A.

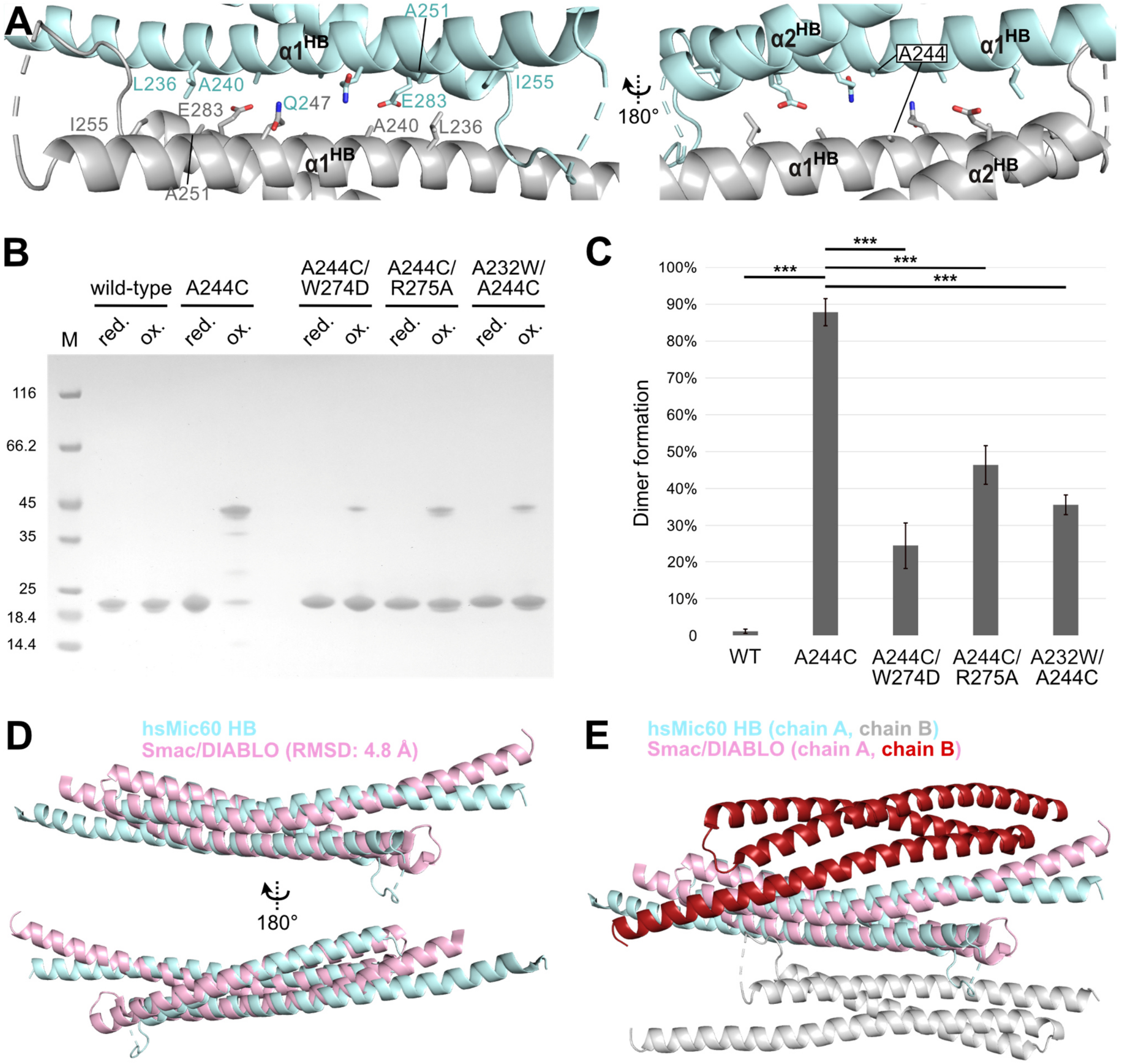


**Supplementary Figure 3: Mic60 HB dimerization is captured by the artificial cysteine bridge A244C and differs from structurally homologous Smac/DIABLO**

**A** Close-up view of the dimerization interface of Mic60 HB, which centres around an antiparallel hydrophobic patch of $\alpha 1^{HB}$ involving residue A244, used in designing artificial cysteine variants of Mic60 HB. **B** Independent replicate experiment of Figure 1D: Non-reducing SDS PAGE of Mic60 HB cysteine mutants before and after $CuSO_4$ oxidation (from left to right: wt, A244C, A244C/W274D, A244C/R275A, A232W/A244C). Lower bands correspond to a monomer, upper bands correspond to a dimer formed through the artificial cysteine bridge. **C** Dimer formation quantification after $CuSO_4$ oxidation for Mic60 HB wild-type and cysteine mutants using non-reducing SDS PAGE (see Figure 1D). Bars indicate the standard deviation from triplicate gels. ***$P \leq 0.005$, calculated by t-test from three replicates. **D** Smac/DIABLO (PDB ID: 1FEW) was identified as a homologous structure to the Mic60 HB using Foldseek (van Kempen et al., 2024) (RMSD = 4.8 Å, Smac/DIABLO monomer coloured pink). **E** Mic60 HB and Smac/DIABLO deploy different dimerization interfaces (Mic60 HB dimer coloured in cyan and grey, Smac/DIABLO dimer coloured in pink and red).

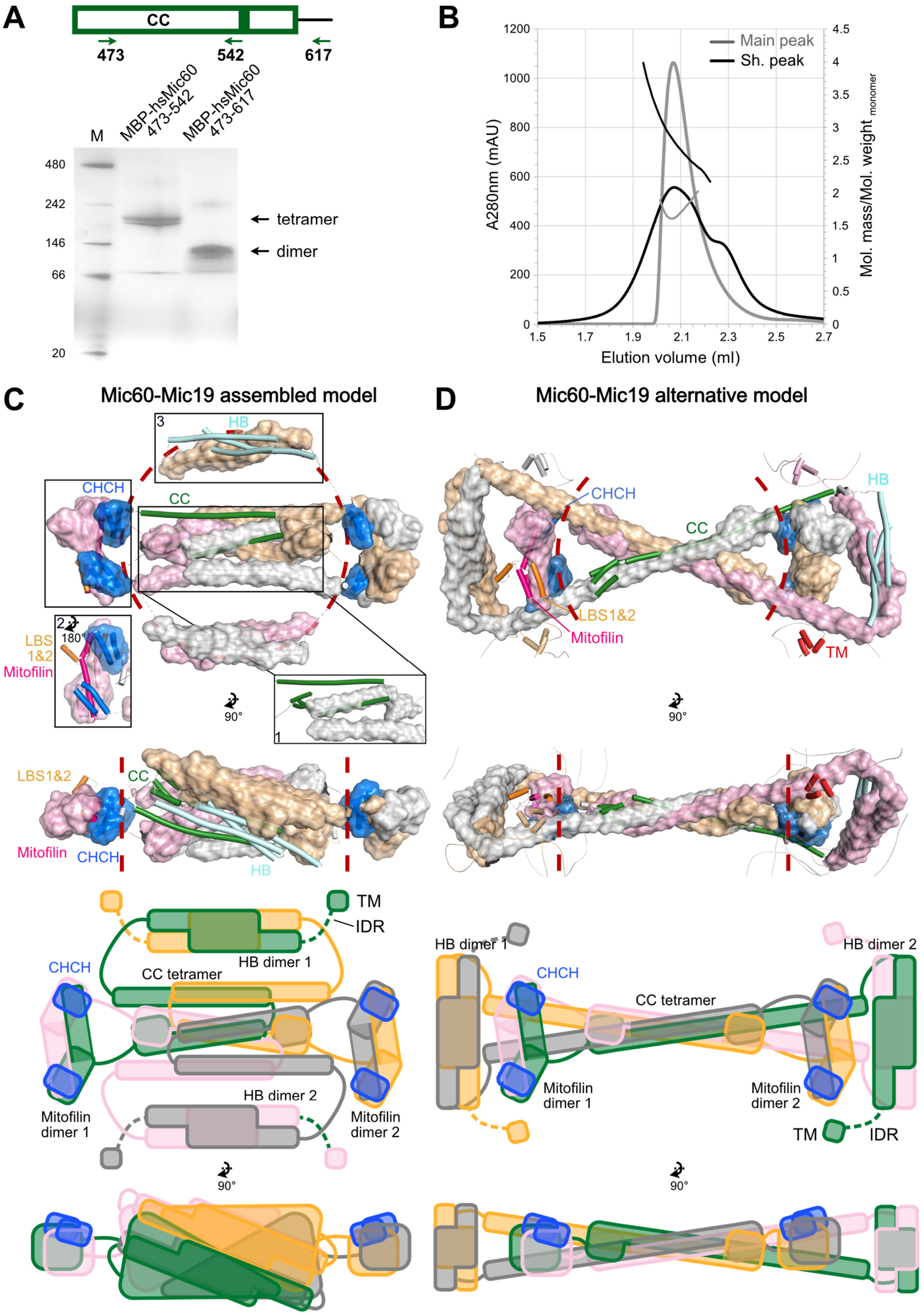

A
CC
473
542
617
M
MBP-hsMic60 473-542
MBP-hsMic60 473-617
480
242
146
66
20
tetramer
dimer
B
Main peak
Sh. peak
A280nm (mAU)
Mol. mass/Mol. weight monomer
Elution volume (ml)
C
Mic60-Mic19 assembled model
D
Mic60-Mic19 alternative model
CHCH
CC
HB
LBS 1&2
Mitofilin
LBS1&2
TM
IDR
HB dimer 1
HB dimer 2
CC tetramer
Mitofilin dimer 1
Mitofilin dimer 2
90°
180°


1249

42

**Supplementary Figure 4: hsMic60 can form tetramers similarly to *Lachancea thermotolerans* Mic60, leading to two possible Mic60-Mic19 complex assemblies**

**A** Graphic illustration of the two hsMic60 CC constructs: CC_1 hsMic60 473-542 and CC_2 hsMic60 473-617. BN-PAGE analysis shows the truncated CC_1 construct can form tetramers while CC_2 can form only dimers. **B** SEC-RALS profile of CC_1. The protein was separated during purification into an earlier shoulder peak (grey) and a main peak (black) and the peaks were pooled separately. They were analysed again via SEC-RALS and eluted in a dimer-tetramer equilibrium and as a dimer, respectively. **C** The main homology-based model of the human Mic60-Mic19 complex (above) along with a graphical cartoon (below) as shown in Figure 2A. **D** An alternative conformation of the model of the human Mic60-Mic19 complex. The model is coloured as in C and fulfils the same structural information as the first model. Here, the two dimeric HB modules are assembled by the same Mic60 moieties that dimerize through their mitofilin domains, resulting in a more extended conformation.

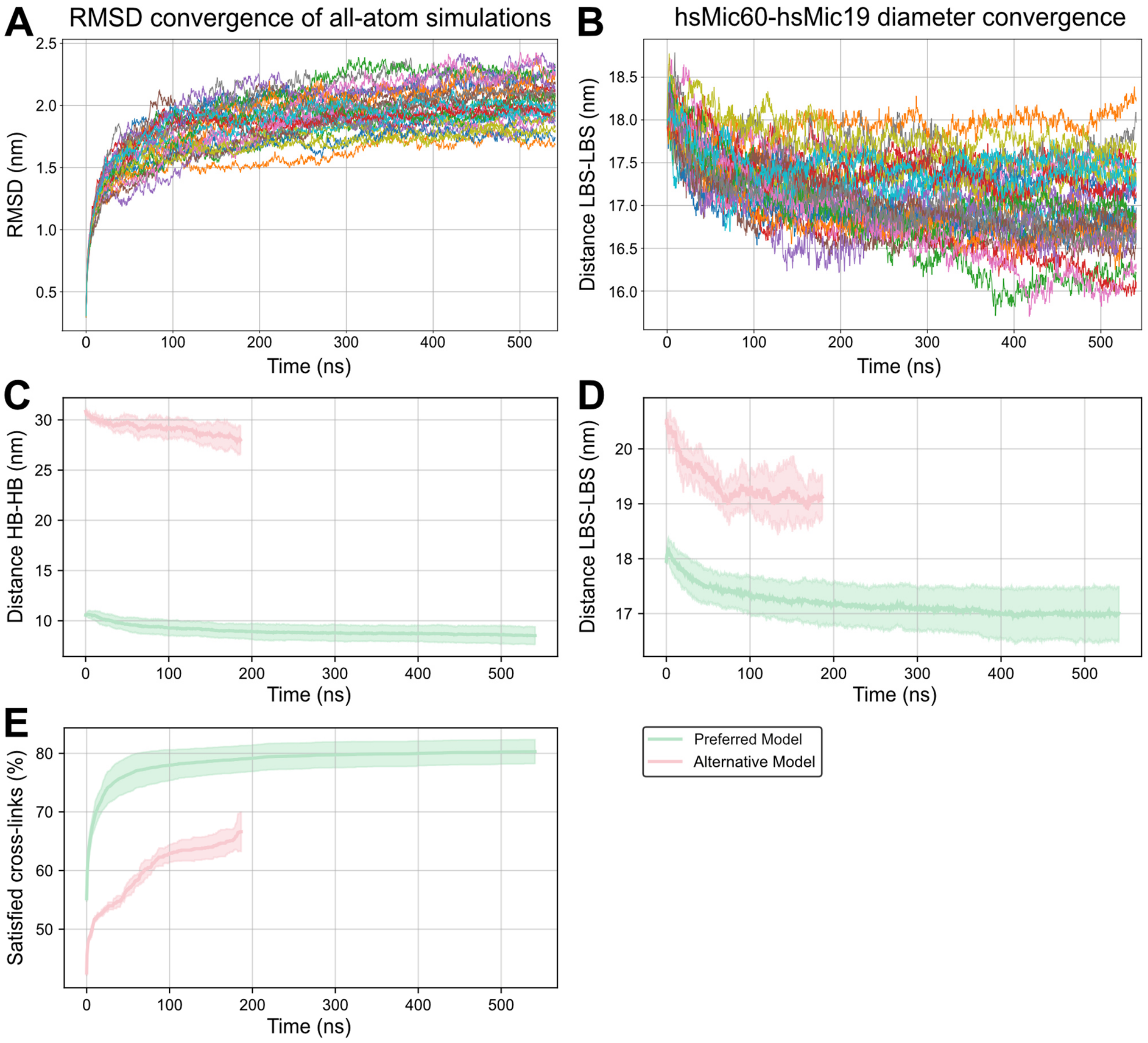


**Supplementary Figure 5: Equilibration graphs (RMSD, HB-HB and LBS-LBS distance) and cross-link validation favour the main model of the human Mic60-Mic19 complex over the alternative model**
**A** RMSD of each of the 30 all-atom simulations is shown over the time course of the simulation, resulting in the average and standard deviation of Figure 2B. **B** LBS-LBS distance for each of the 30 all-atom simulations, resulting in the average and standard deviation of Figure 2C. **C, D** Diameter convergence of the preferred and alternative human Mic60-Mic19 complex models during the all-atom simulations, prior to the attachment to the catenoid, plotted for the HB-HB distance (C) and the LBS-LBS distance (D). Line represents the mean and shading represents the standard deviation across three independent trajectories. **E** The percentage of cross-links satisfied cumulatively across the frames of the all-atom simulation of the preferred and the alternative models for all possible cross-links. Rather than considering all independent simulations in parallel as in Figure 3B, the mean and standard deviation over the independent runs was plotted to account for the different number of trajectories, resulting in a lower satisfaction rate compared to Figure 3B. Line represents the mean and shading represents the standard deviation across three independent trajectories for the alternative model and thirty independent trajectories for the preferred model.

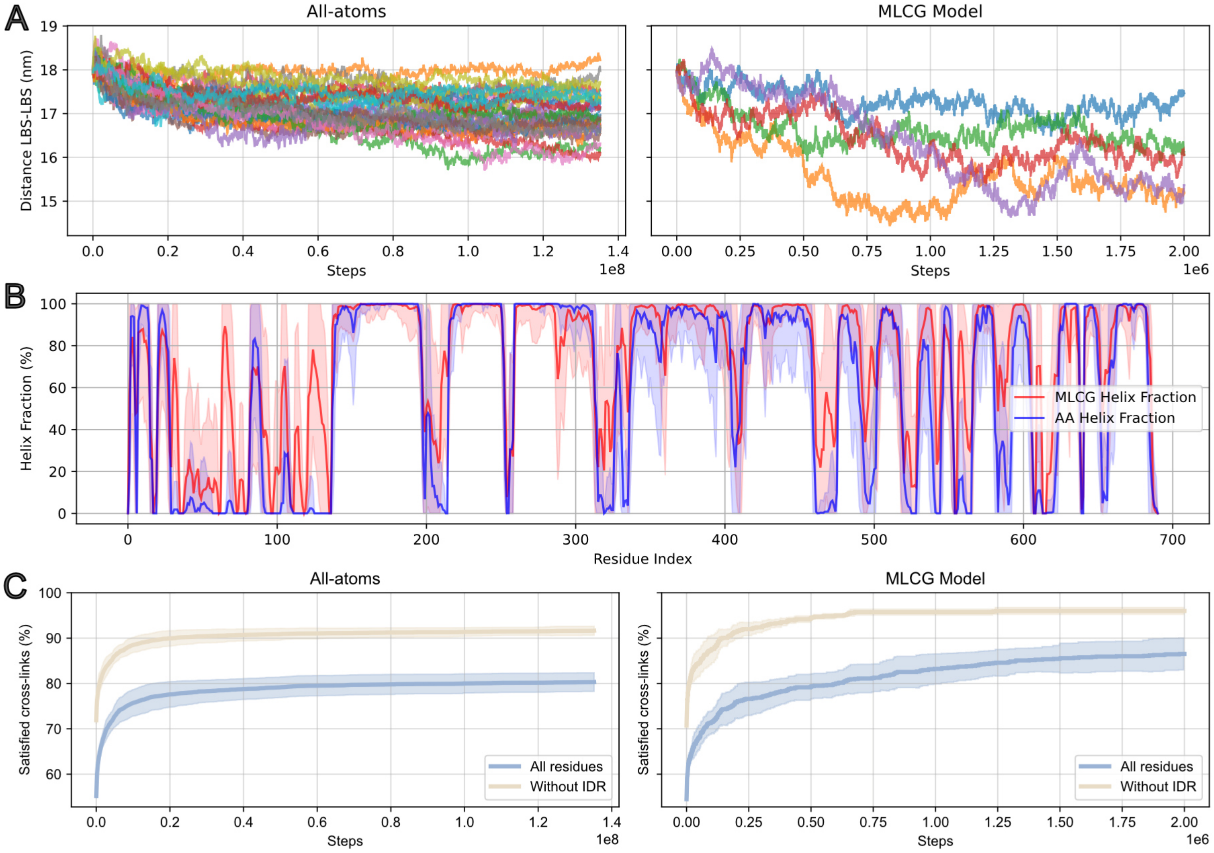


**Supplementary Figure 6: Machine-learned coarse-grained (MLCG) model simulations**

**A** Diameter convergence (plotted as LBS–LBS distance) across the steps of the simulation, comparing the all-atom and MLCG simulations, exhibiting a more pronounced structural compression and greater variability across independent trajectories for the MLCG (right) compared to the all-atom model (left). **B** Helical content per residue, determined using coarse-grained DSSP in MDTraj (McGibbon et al., 2015) and shown as the mean and standard deviation over the simulation, highlights the correspondence between the all-atom and MLCG models. **C** The percentage of cross-links satisfied across the steps of the all-atom or MLCG simulations was calculated as in Supplementary Figure 5E. Line represents the mean and shading represents the standard deviation across thirty independent trajectories for the all-atom simulations and six independent simulations for the MLCG model.

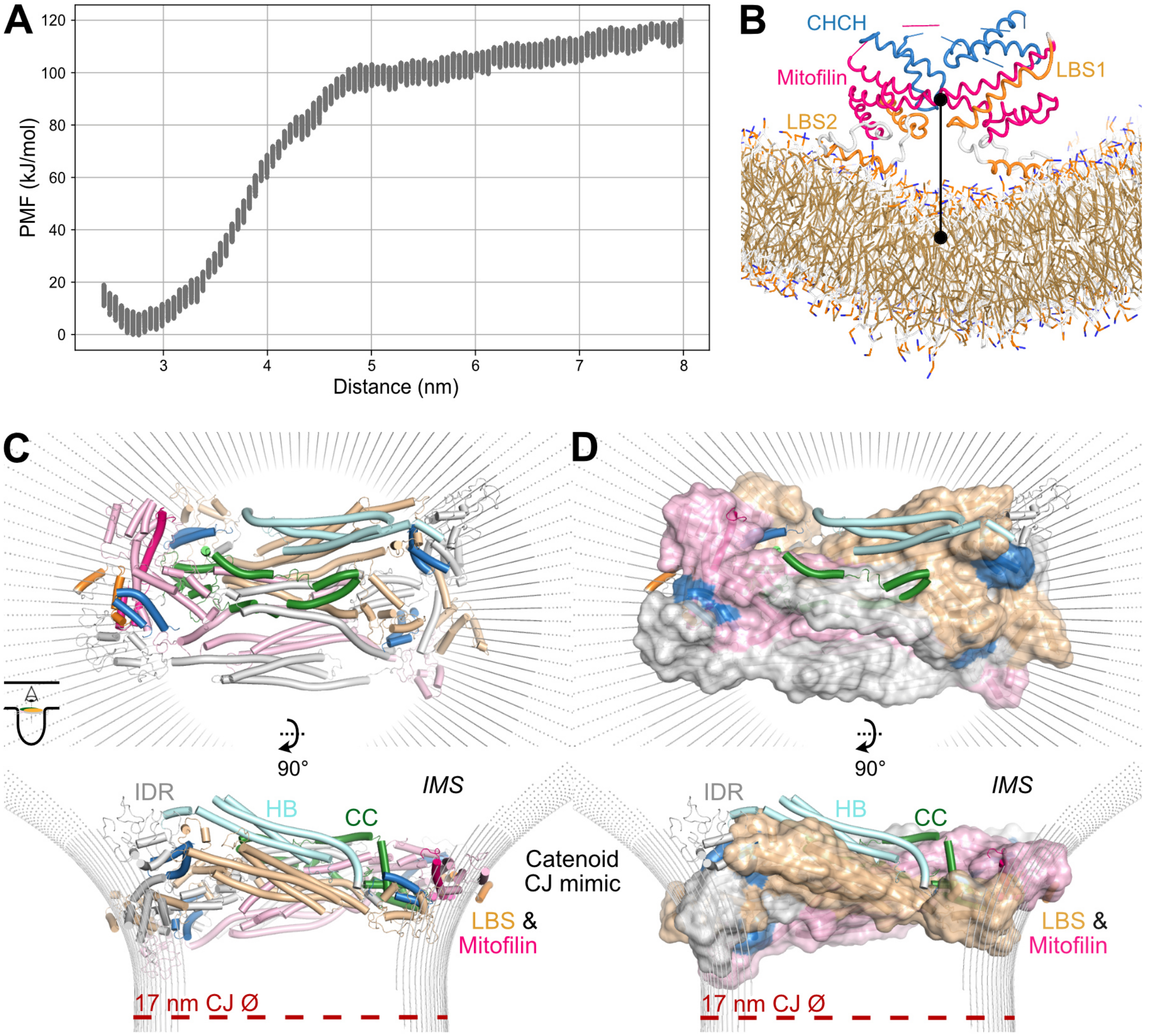


**Supplementary Figure 7: IMM attachment experiments and all-atom MD simulations for human Mic60-Mic19 on a CJ mimic**

**A** Potential-mean-force (PMF) plot of the interaction of the C-terminal mitofilin-CHCH lobe to an IMM mimic. A release of approximately 100 kJ/mol is observed upon binding. **B** Cartoon illustration of the interaction of the C-terminal mitofilin domain dimer (containing the mitofilin and the CHCH domains) to an IMM mimic. Black circles indicate the centre of each object, while a black bar indicates their distance, as plotted in A. **C, D** Final snapshots of the all-atom MD simulations after attachment to the catenoid, with the model represented as cylinders (C) or cylinders and surface (D). The base diameter of the catenoid, indicated in a red dashed line, is 17 nm. Note the compression of the IDRs in comparison to the Cα structure-based MD simulations shown in Figure 2D.

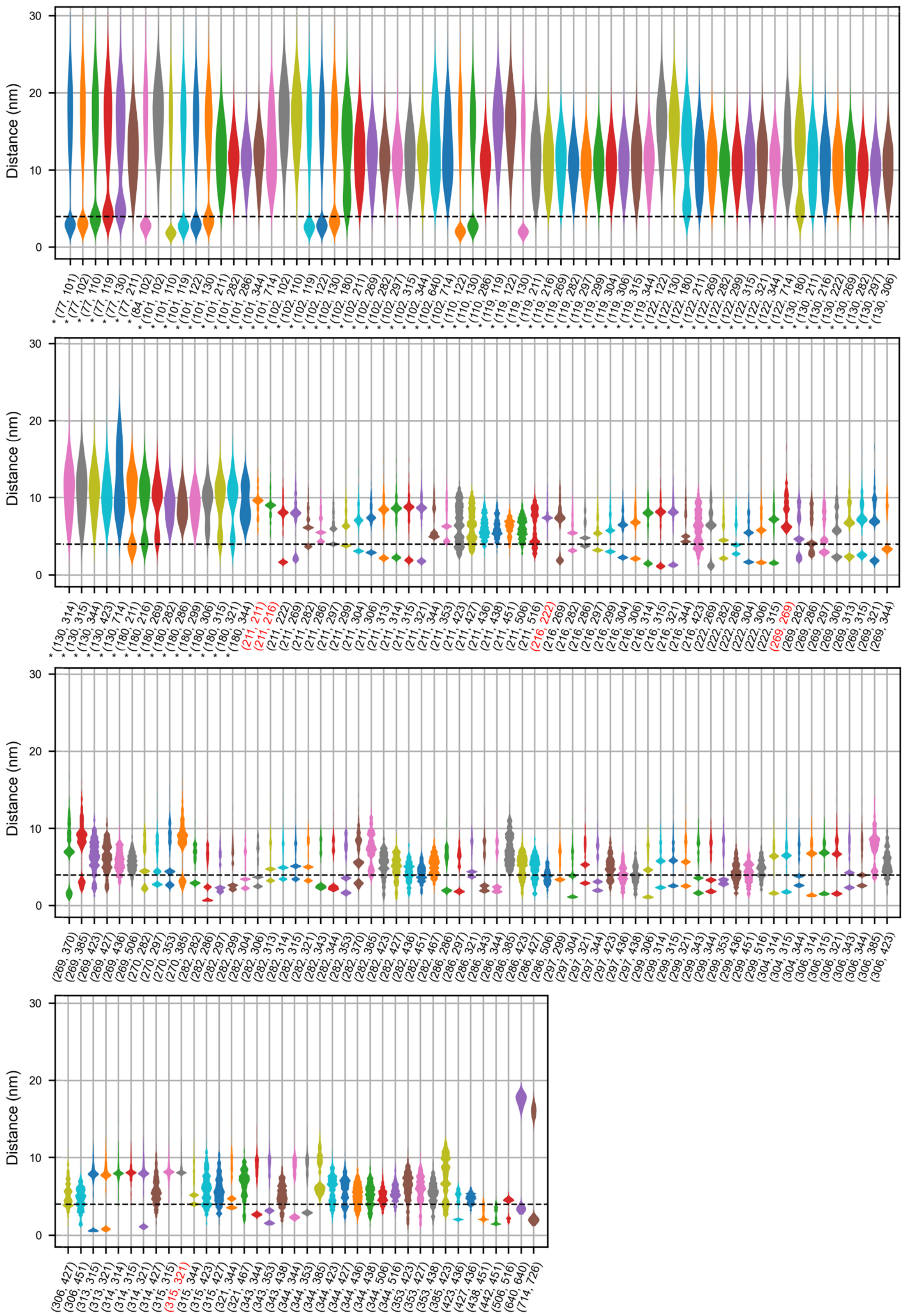
Distance (nm)

**Supplementary Figure 8: hsMic60 cross-linking dataset for MD validation**

Violin plot of all 236 cross-links present in the dataset employing the enrichable and MS-cleavable cross-linker DSBSO (Zhu et al., 2024). The cross-link distance is plotted for each residue pair, including self-cross-links, across all Cα structure-based simulation frames. The threshold for validity was set at 4 nm (40 Å). Cross-links involving the IDR are marked with an asterisk. Cross-links that are not valid in any of the frames of the simulations are marked red.

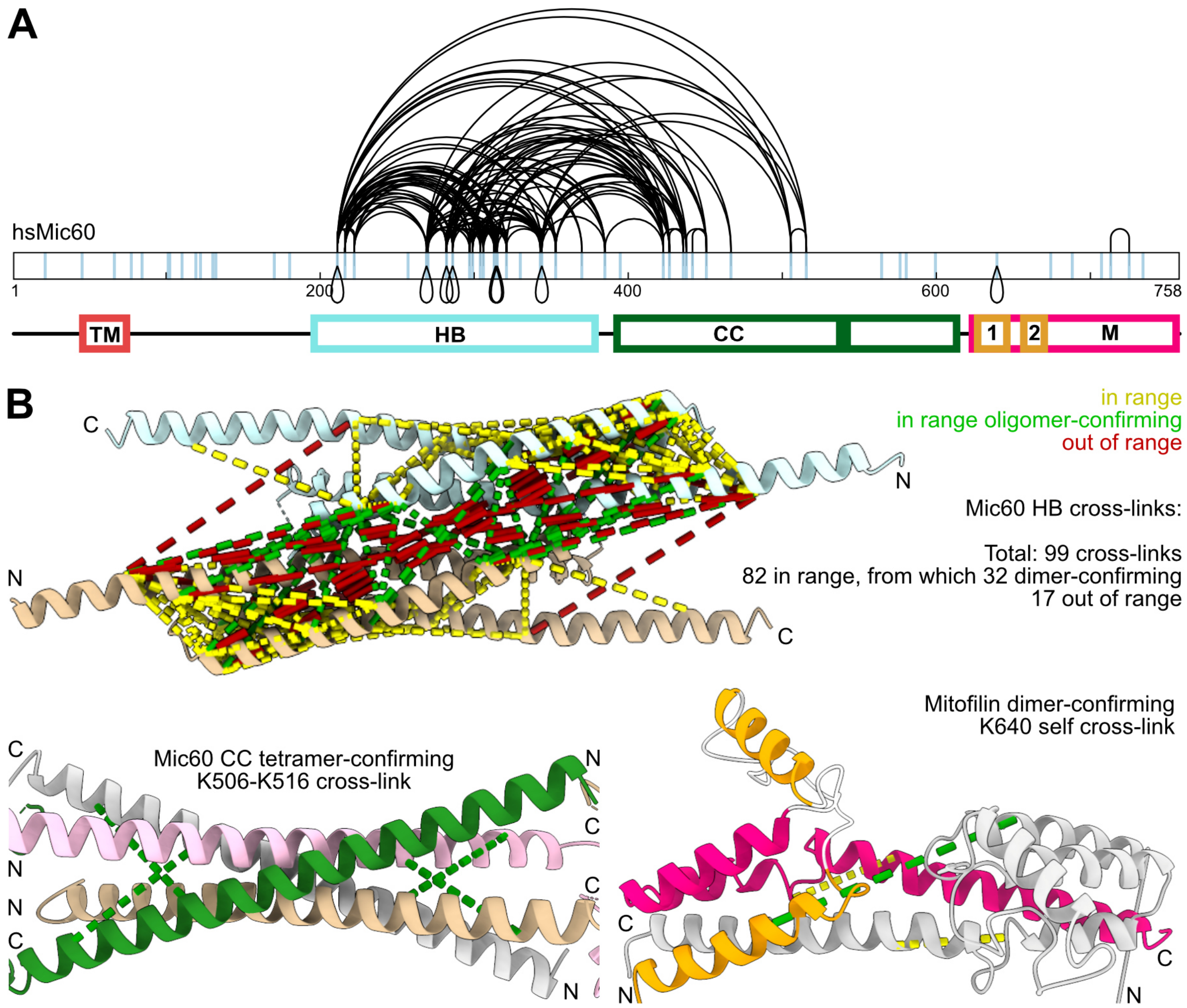


**Supplementary Figure 9: *In organello* cross-linking as validation of the homology model prior to MD simulations**

**A** Visualisation of the DSBSO mitochondrial cross-linking dataset on the Mic60 sequence as in Figure 3A, excluding the cross-links involving the IDR, generated with xiNET (Combe et al., 2015). Blue bars across the protein sequence indicate the positions of lysines. **B** Analysis of the intra-domain cross-links of Mic60 before MD simulations. Cross-links under 40 Å are in range and coloured in yellow, with those in range cross-links that confirm domain homo-oligomerization coloured in green, and out of range cross-links in red. For the Mic60 HB dimer 99 cross-links were detected (82 of them in range, with 32 of those confirming a HB dimer), for the Mic60 CC tetramer one cross-link was detected and confirmed a homo-tetrameric assembly, and for the mitofilin dimer two cross-links were detected, one of which corresponded to a dimeric conformation.

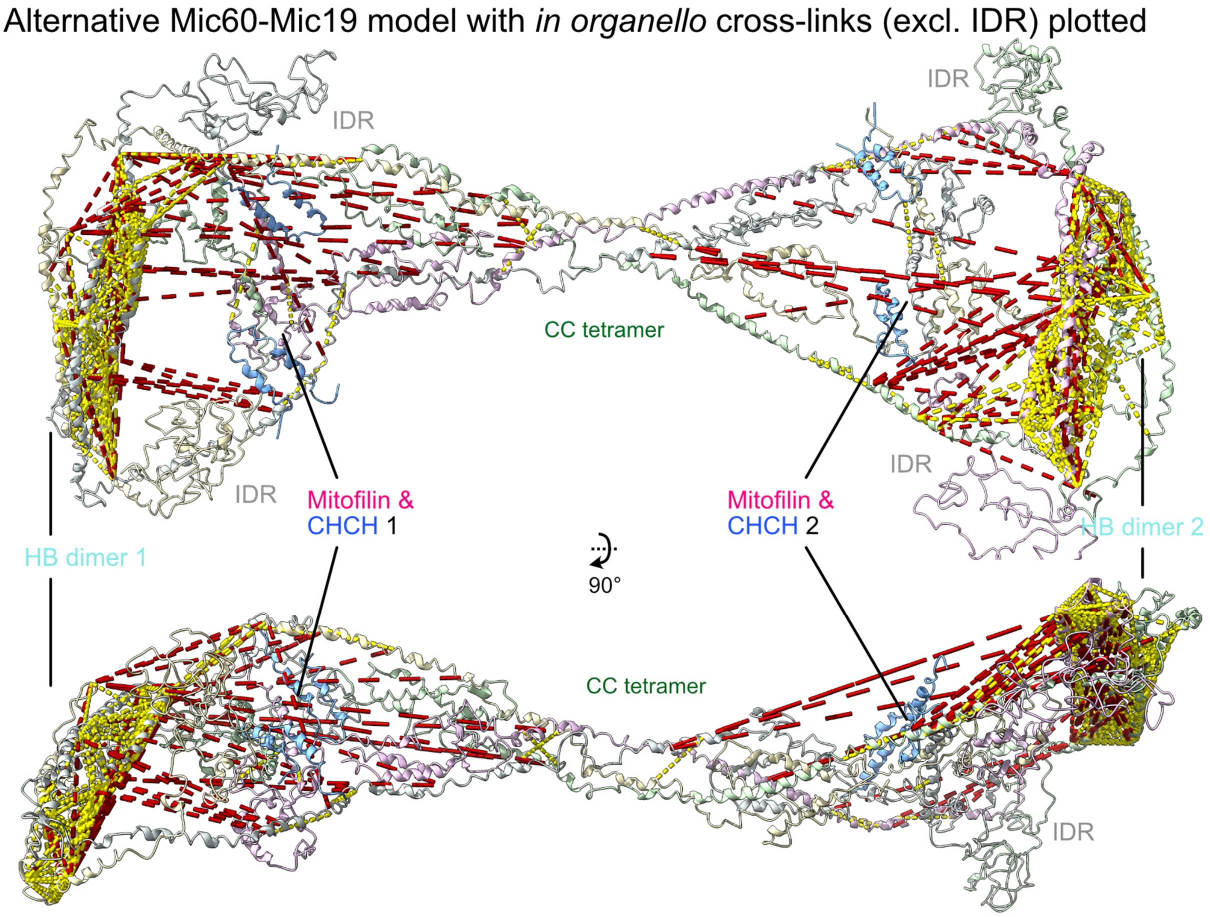


**Supplementary Figure 10: Cross-linking comparison between the two homology-based models**
The cross-linking dataset (excluding IDR cross-links for ease of illustration) plotted on the alternative homology-based Mic60-Mic19 model after the all-atom simulations (yellow for in range, red for out of range).

**Supplementary Table 1: Data collection and refinement statistics (molecular replacement)**

| | Mic60 HB (PDB ID: 9QWR) |
|---|---|
| **Data collection** | |
| Beamline | BESSY 14.1 |
| Wavelength (Å) | 0.9184 |
| Space group | $P4_12_12$ (92) |
| Cell dimensions | |
| *a*, *b*, *c* (Å) | 76.12 76.12 168.41 |
| $\alpha, \beta, \gamma$ (°) | 90.0 90.0 90.0 |
| Resolution (Å)* | 33.16 - 2.78 (2.88 - 2.78) |
| $R_{merge}$ (%) | 13.7 (324.2) |
| CC(1/2) (%) | 99.9 (30.1) |
| $I / \sigma I$ | 12.48 (0.67) |
| Completeness (%) | 99.6 (97.4) |
| Redundancy | 11.4 (11.5) |
| **Refinement** | |
| $R_{work}$ / $R_{free}$ (%) | 24.5 / 28.3 |
| Reflections | 13,089 (2,028) |
| No. atoms | |
| Protein | 2,542 |
| Water | 1 |
| B-factor | |
| Protein | 94.35 |
| R.m.s. deviations | |
| Bond lengths (Å) | 0.003 |
| Bond angles (°) | 0.48 |
| Ramachandran Plot | |
| Outliers (%) | 0.00 |
| Allowed (%) | 0.62 |
| Favoured (%) | 99.38 |
| Rotamer outliers (%)$ | 0.75 |
| Clashscore | 5.23 |

*Values in parentheses are for highest-resolution shell. Correlation coefficient between two independent subsets (CC1/2) according to Karplus and Diederichs (2012). Model geometry according to Molprobity (Williams et al., 2018).

**Supplementary Video 1: The human Mic60-Mic19 complex as a diffusion barrier in CJs**

Video of all simulation frames of the Cα-based diffusion experiments of the human Mic60-Mic19 model in the catenoid. 200 spheres, here with the radius of 1.5 nm, were allowed to pass through the junction. There were no repulsion forces between the spheres, only between the spheres and the complex or the catenoid, allowing them to be simulated simultaneously.
